# Pathway Representation via Intrinsic Structural Medoids (PRISM): A Structural Mapping Approach to Clustering Molecular Pathways

**DOI:** 10.64898/2026.05.16.725628

**Authors:** Jherome Brylle Woody Santos, Jeremy M. G. Leung, Lillian T. Chong, Ramón Alain Miranda-Quintana

## Abstract

We present Pathway Representation via Intrinsic Structural Medoids (PRISM), a state-aware framework for clustering pathways from molecular dynamics simulations of biomolecular transitions. In PRISM, each pathway is mapped to a small set of structural medoids obtained via a deterministic *k*-means clustering scheme. Pairwise pathway dissimilarities are computed using a weighted average Hausdorff distance between these representative sets, effectively capturing mean nearest-neighbor structural deviations while reducing sensitivity to outliers. Hierarchical agglomerative clustering of the resulting dissimilarity matrix defines pathway families. We evaluate PRISM across three biomolecular transitions of increasing complexity: alanine dipeptide C7_eq_ → C7_ax_ isomerization, adenylate kinase opening, and HIF-2α PAS-B ligand unbinding. PRISM consistently yields robust cluster assignments, with medoids faithfully representing distinct conformational states. By combining a state-based description with robust geometric dissimilarities, PRISM provides a scalable framework for organizing complex transition pathways.

## 1. INTRODUCTION

Advances in both computing hardware and software have greatly extended the timescales accessible to atomistic molecular dynamics simulations.^1–5^ In addition, path sampling strategies such as transition path sampling,^6^ milestoning,^7^ and the weighted ensemble (WE) strategy^8^ are designed to efficiently generate ensembles of pathways for barrier-crossing processes. While these path ensembles are rich in mechanistic information, they also present a substantial analysis challenge: the individual pathways are numerous, heterogeneous in length, and often explore overlapping yet distinct regions of the conformational space.^9–11^ A central goal, therefore, is to organize these data into pathway families that capture dominant mechanisms while also revealing less probable but functionally relevant alternatives.

Several pathway clustering approaches have been developed to address this goal. One broad class relies on geometric distances between entire trajectories, such as Hausdorff or Fréchet metrics computed from pairwise structural distance between pathway frames, as in Path Similarity Analysis (PSA).^10^ These methods compare pathways as continuous curves and have been proven useful for quantifying large-scale mechanistic differences. Moreover, the linguistic-based strategy, Linguistics Pathway Analysis of Trajectories with Hierarchical clustering (LPATH),^11^ discretizes configuration space into states and represents each pathway as a text string of states visited, enabling rapid pairwise comparisons through a string-similarity score followed by hierarchical clustering. More recently, deep learning approaches have been proposed, in which trajectories are embedded into a learned low-dimensional latent-space representation prior to clustering.^12–14^ Across these frameworks, the design of a pathway similarity measure remains the key ingredient: it must distinguish among distinct routes while remaining computationally efficient.

Our recent work, Sampling Hierarchical Intrinsic *N*-ary Ensembles (SHINE),^9^ introduced a different approach to pathway clustering based on *n*-ary similarity measures. Rather than relying solely on pairwise frame distances between trajectories, SHINE constructs a pairwise pathway dissimilarity matrix using set-based measures defined over the union of frames across all pathways. SHINE incorporates several pathway similarity metrics, including intra- and inter-dissimilarities, as well as semi-sum, minimum, and maximum criteria. Owing to the *n*-ary formulation (i.e., the simultaneous evaluation of collective similarity across *n* inputs), these quantities can be computed in linear time with respect to the number of frames.^9,15–24^ This framework enables deterministic, many-to-many clustering of pathways and has been shown to recover pathway classes for conformational transitions in alanine dipeptide and adenylate kinase.

Despite these advantages, several limitations remain when pathway dissimilarities are defined directly on full frame ensembles. First, the mean-squared deviation (MSD) and related metrics for dissimilarity treat each pathway as an amorphous cloud of structures. While this is advantageous for efficiency, it obscures how trajectories populate specific regions of conformational space, making it harder to connect pathway families back to interpretable structural states.^25^ Second, Hausdorff-type distances, when defined on all frames, are dominated by worst-case outliers: a single excursion can cause two otherwise similar pathways to appear maximally dissimilar. This sensitivity is particularly problematic for noisy or highly flexible systems, in which transient fluctuations are common. Third, computing ensemble-level distances between full trajectories can become costly for large path ensembles (e.g., beyond thousands of pathways), motivating either aggressive sampling or additional approximations.

In parallel, our recent work on clustering individual molecular dynamics (MD) simulation trajectories has demonstrated the value of combining *k*-means pre-clustering within hierarchical agglomerative clustering through *n*-ary similarity metrics. The Hierarchical Extended Linkage Method (HELM)^22^ framework showed that the deterministic *k*-means *N*-Ary Natural Initiation (NANI)^26,27^ and MSD-based linkage can dramatically reduce the cost of hierarchical clustering while preserving the detailed hierarchical structure. These developments suggest that similar hybrid strategies could be leveraged at the pathway level, where we would like to retain the mechanistic focus on path clustering while benefiting from the interpretability and efficiency of state-based representations.

Here, we introduce Pathway Representation via Intrinsic Structural Medoids (PRISM), a new method for clustering molecular pathways that builds on the ideas of SHINE while adopting a distinct state-aware design. Rather than comparing the full set of conformations in each pathway, PRISM first maps every pathway onto a compact set of representative conformations: structural medoids obtained from *k*-means NANI clustering in a chosen feature space. These medoids serve as intrinsic “states” for each trajectory. Pairwise pathway dissimilarities are then quantified using a weighted average Hausdorff distance between the representative sets. This metric measures the mean nearest-neighbor mismatch between medoid sets in both directions, providing a robust measure of pathway overlap that downweights isolated outliers and emphasizes the common structural neighborhoods explored along each route.

PRISM offers several advantages relative to existing pathway-clustering frameworks. By working with medoids, it establishes an explicit bridge between pathway families and structural states, facilitating direct structural interpretation of each cluster. The use of representative sets also reduces the effective size of each trajectory, which in turn lowers the cost of building the dissimilarity matrix, especially for very long or densely sampled pathways. Moreover, the weighted average Hausdorff formulation retains a clear geometric meaning while avoiding some of the disadvantages of the classical Hausdorff distance for biomolecular ensembles.

In this work, we develop the theoretical basis of PRISM and explore three medoid-construction strategies that differ on how they balance global and pathway-specific information. We then apply the method to three benchmark processes: the C7_eq_ to C7_ax_ transition in alanine dipeptide,^9,11^ the closed-to-open transition in adenylate kinase,^10,28,29^ and the seconds-timescale unbinding of a drug-like ligand from a completely buried cavity of the HIF-2α PAS-B domain.^30^ Across these processes, we assess PRISM’s ability to recover known mechanistic classes, its sensitivity to method parameters, and its computational efficiency relative to the original SHINE intra scheme and its sampling variants. Overall, these results establish PRISM as a state-aware, structurally interpretable alternative for clustering molecular pathways, complementing existing approaches and extending MDANCE,^21^ our MD simulation analysis and clustering package, for pathway analysis in the exascale era of molecular simulations.

## 2. THEORY

### Pathway-level dissimilarities in PRISM

Let T = {*T*_1,_ …, *T*_*p*_}be a collection of pathways. Each pathway *T*_*i*_ is a trajectory of *n*_*i*_ frames embedded in a common feature space ℝ^*d*^, as defined in Eq. (1),

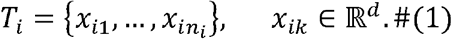

In SHINE, we quantify pathway-pathway relationships by constructing a dissimilarity matrix over T and applying hierarchical agglomerative clustering to this matrix. The original framework defines various dissimilarity formulations based on *n*-ary metrics evaluated on *T*_*i*_, *T*_*j*_, and their union *T*_*i*_ ⋃ *T*_*j*_. These include: intra, the MSD of the full union; inter, the MSD of points between *T*_*i*_ and *T*_*j*_ (analogous to average linkage); semi-sum, the union MSD minus the average of the individual trajectory MSDs; and min/max, the union MSD minus the minimum/maximum of the individual trajectory MSDs. The classical Hausdorff distance is also supported. These formulations are expressed in terms of ensemble-level comparisons between frames.

PRISM is a method that implements a new pathway-pathway dissimilarity definition based on the weighted average Hausdorff distance, as applied to compact sets of representative frames extracted from each trajectory.

### Weighted average Hausdorff distance

Let 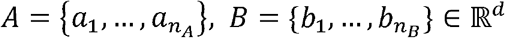 be finite sets of representative frames associated with two pathways with *n*_*i*_ number of frames. For each point *a*_*p*_ ϵ *A*, its nearest-neighbor distance to *B* is defined in Eq. (2) as

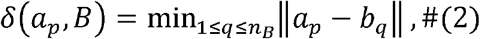

and similarly, for each *b*_*q*_ ϵ *B*, the nearest-neighbor distance to A is given by Eq. (3),

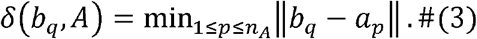

The directed average distances from *A* to *B* and from *A* to *B* read as in Eqs. (4) and (5),

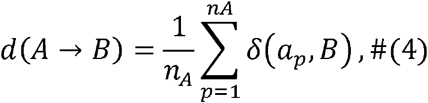

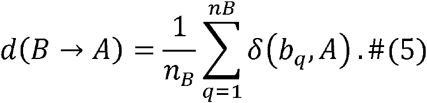

We then define the weighted average Hausdorff distance between A and B as Eq. (6),

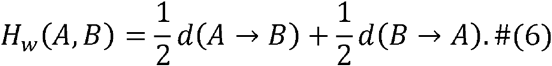

By construction, *H*_*w*_ (*A,B*) is non-negative, symmetric, and equals zero when two representative sets are identical. Although the distance generally does not satisfy the triangle inequality and is, therefore, not a true metric in the strict mathematical sense,^10^ it is sufficient for our application, as hierarchical clustering in PRISM (and in SHINE) relies on relative pairwise dissimilarities rather than metric space embeddings. Crucially, this formulation preserves a highly interpretable geometric meaning that is physically superior to the classical Hausdorff distance for biomolecular ensembles. Rather than defining similarity by the worst-case outlier, *H*_*w*_ measures the mean nearest-neighbor mismatch in both directions, providing a robust assessment of structural overlap that effectively filters out transient noise.

### Construction of representative sets

The distance *H*_*w*_(*A,B*) is formulated for arbitrary sets of points from both *A* and *B*. To apply it to full trajectories, each pathway *T*_*i*_ is mapped to a finite representative set *A*_*i*_. We construct the representatives from medoids of clusters in the feature space.

Here, we consider three options for building *A*_*i*_, which differ in whether clustering is performed globally across all pathways, locally per pathway, or in a two-stage fashion that combines both. In all three cases, clustering is performed with the *k*-means NANI algorithm, a *k*-means variant, which provides a robust and efficient determination of initial cluster centers. *N*-ary calculations also allow us to deterministically and efficiently estimate the medoid, which we define as the frame with maximal complementary similarity to the rest of the frames in its cluster, ensuring that the representative is the most central conformation within the subset.

#### Option 1: Global medoid sharing

In the first option, all frames from all T pathways are pooled into a single dataset, as shown in Eq. (7),

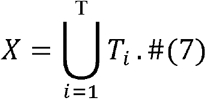

*X* is then partitioned into *k* clusters. A medoid is identified for each cluster.

Each pathway *T*_*i*_ is now represented by the subset of global medoids whose clusters contain at least one frame from *T*_*i*_. Let {*m*_1_,…,*m_k_*}, be the medoids and let *C*_*c*_ ⊂ *X* denote the cluster associated with *m*_*c*_. The representative set for pathway *i* is expressed in Eq. (8) as

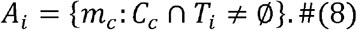

In this construction, the assumption is that pathways that explore similar regions of conformational space will share many global medoids, whereas pathways that are geometrically distinct will share few or none. The average Hausdorff distance between *Ai* and *A*_*j*_ therefore reflects how similarly each pathway occupies the global landscape of conformations.

#### Option 2: Independent medoids per pathway

In the second option, each pathway is grouped independently. For a given trajectory *Ti*, its frames are clustered into *k* clusters, and their medoids are computed. This yields a pathway-specific set of medoids where *k*_*i*_ ≤ *k*, as defined in Eq. (9),

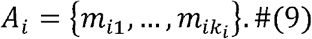

Here, representatives are not shared across pathways; medoids of *Ti* and *Tj* are distinct even if the underlying frames reside in similar regions of conformational space. The distance *Hw*(*A_i_,A_j_*)is steered by how these local, pathway-specific representatives can be combined. This option emphasizes the internal structure of each trajectory and therefore can be more sensitive to differences in how pathways traverse similar regions.

#### Option 3: Two-stage medoid refinement

In the third and last option, we combine the ideas from the first two options in a two-stage scheme. In the first step, each pathway *T_i_* is clustered independently as in Option 2, yielding a local initial set of medoids 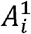. These representatives from all pathways are collected into a pooled set *M*, as shown in Eq. (10),

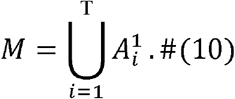

In the second stage, *M* is clustered into *k*_final_ clusters, and a medoid is computed for each of these. These final medoids form a refined global dictionary 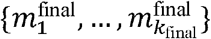. A pathway *T*_*i*_ is then represented by those final medoids whose clusters contain at least one of its local medoids. If 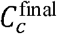 denotes the set of local medoids assigned to the c-th cluster, the representative set for pathway *i* is expressed in Eq. (11) as

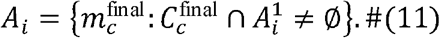

This allows each pathway to first compress its internal variability into a number of local medoids and then align these local summaries through a global clustering step. It thus provides a compromise between the purely global perspective of Option 1 and the purely local perspective of Option 2.

Once representative *A*_*i*_ sets have been defined by one of the three options above, we can proceed with building the pathway-pathway dissimilarity matrix with entries *D*_*ij*_=*H*_*w*_(*A*_*i*_,*A*_*j*_)and apply hierarchical clustering (Fig. 1), following the procedure of the original SHINE framework.

**Figure 1.**
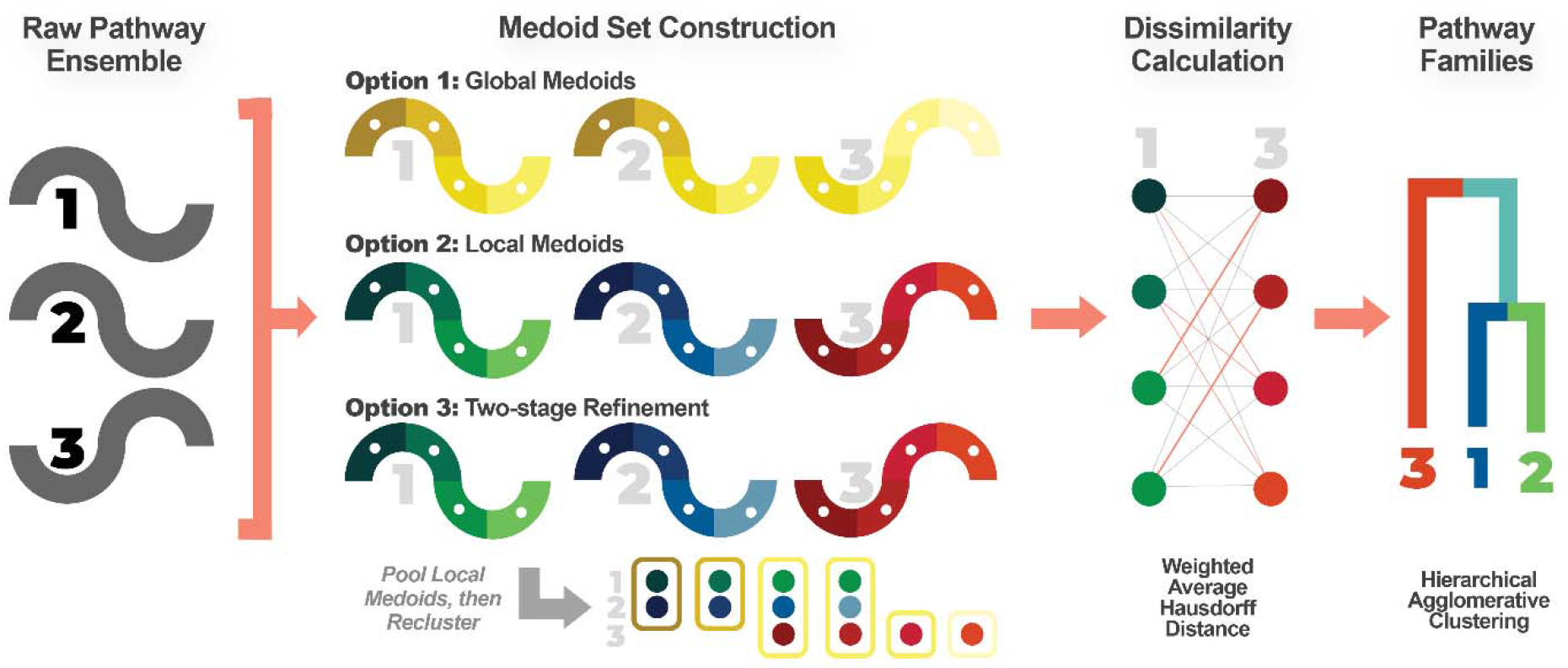
Overview of the PRISM workflow. A raw ensemble of molecular transition pathways is mapped to compact sets of medoids via one of three construction strategies. In Option 1, all pathway frames are pooled and clustered jointly, and each pathway is represented by the subset of global medoids whose clusters it occupies. In Option 2, each pathway is clustered independently, yielding a pathway-specific medoid set. In Option 3, local medoids are first constructed per pathway, pooled, and then re-clustered globally to produce a refined shared dictionary. Pairwise pathway dissimilarities are computed using the weighted average Hausdorff distance between medoid sets. The resulting dissimilarity matrix is then passed to hierarchical agglomerative clustering to obtain pathway families.

### Ordering versus occupancy in pathway comparisons

PRISM compares pathways through representative sets of structural medoids, and the weighted average Hausdorff dissimilarity therefore emphasizes structural overlap (occupancy) between pathways rather than the temporal ordering of frames. We intended this design choice to reflect that in many pathway ensembles, mechanistic classes are largely determined by which conformational neighborhoods and intermediates are visited, while short-lived excursions or local fluctuations should not dominate the dissimilarity.

Ordering-aware curve metrics (e.g., Fréchet distance^31^ used in PSA) provide a complementary notion of similarity accounting for the sequential progression along a pathway. Such metrics can be advantageous when pathways visit similar structural regions but differ primarily in the sequence of events or in backtracking behavior. In contrast, PRISM is tailored for robust pathway clustering when the dominant mechanistic modes are encoded in which states are populated and how strongly their representative neighborhoods overlap. Accordingly, PRISM prioritizes structural occupancy as the primary organizing principle of pathway families, while remaining conceptually compatible with ordering-sensitive analyses when traversal sequence is expected to carry additional mechanistic information about the system.

## 3. SIMULATION DETAILS

To evaluate PRISM, we applied the method to the same benchmark processes we used in the original SHINE work:^9^ alanine dipeptide (ALA), adenylate kinase (AdK) and an additional protein-ligand complex consisting of the HIF-2*α* PAS-B domain and small-molecule ligand, THS-017 (HIF-2*α*). Below we summarize the simulation protocols.

### Alanine dipeptide (ALA)

We analyzed an ensemble of 80 successful transition pathways connecting the *C*7_*eq*_ and *C*7_*ax*_ conformational states of alanine dipeptide (Fig. 2). These pathways were obtained from five independent weighted-ensemble (WE) simulations. These runs were performed using WESTPA 2.0 package^32^ with a sampling interval τ 100 ps and a two-dimensional progress coordinate defined by the backbone torsion angles *ϕ* and *ψ*. Binning for the WE resampling was applied only along the *ψ*-dimension, with fixed bins spaced at 20° intervals between 0° to 360°, and a target of eight trajectories per bin.

**Figure 2.**
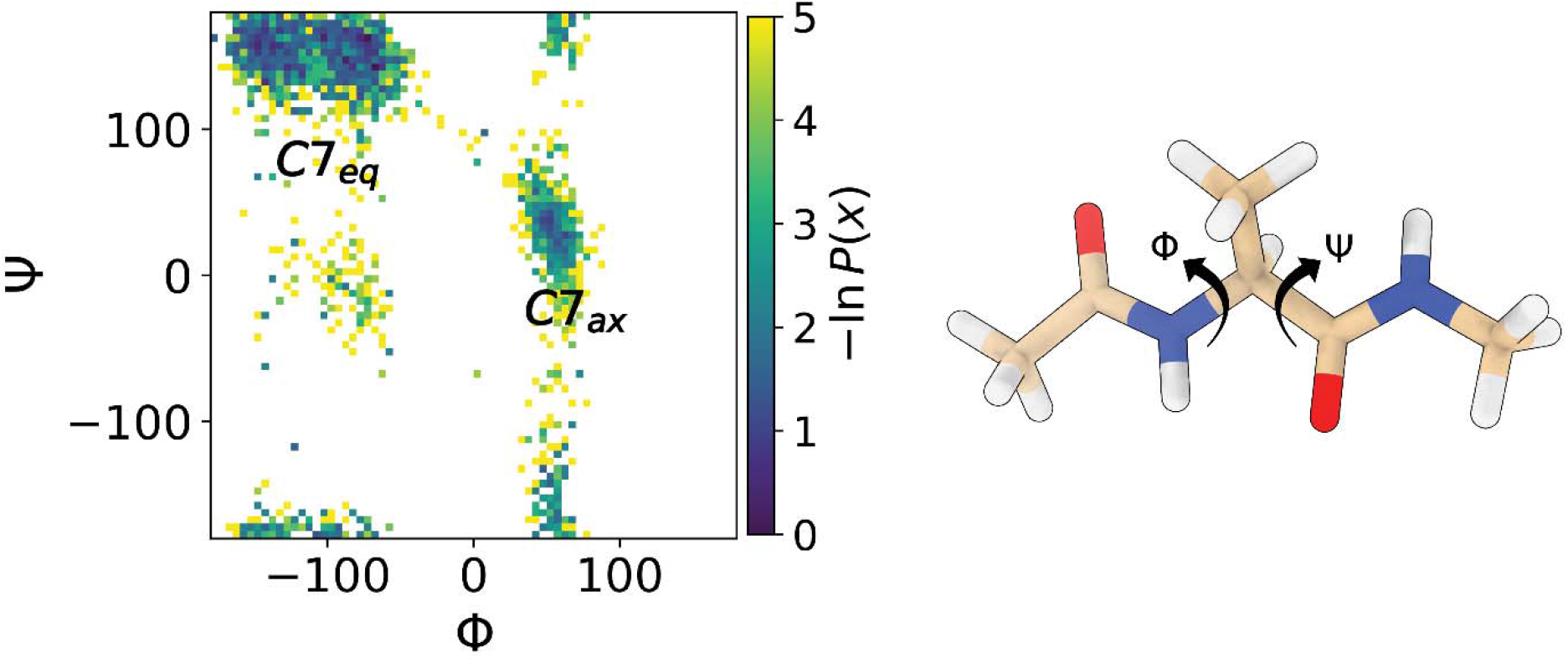
(Left) The 2D free energy surface mapping the 80 successful transition pathways across the backbone torsion angles *ϕ* and *ψ*, showing the distinct *C*7_*eq*_ and C7_*ax*_ basins. (Right) Molecular representation of alanine dipeptide highlighting the specific dihedral angles that drive the transition progress coordinate.

Molecular dynamics for the WE simulations were performed using the Amber 22 package using the Amber ff14SBonlysc^33^ force field in combination with generalized Born implicit solvent (igb = 1).^34^ Hydrogen mass repartitioning was employed to allow a 4 fs integration time step. Across the five WE runs, the aggregate simulation time was 14.6 *µs*, with conformations recorded every 4 ps for subsequent analysis. Each WE simulation was executed on eight CPU cores (3.5 GHz Intel Xeon) and completed in ∼21 hours.

### Adenylate kinase (AdK)

For adenylate kinase (AdK) from *Escherichia coli*, we used a collection of 400 closed-to-open transition pathways provided by Beckstein and co-workers (Fig. 3).^10,28^ Half of these trajectories (200 pathways) were generated using the CHARMM c36b2 engine^35^ with dynamic importance sampling molecular dynamics (DIMS)^29^, a perturbation MD approach that biases random walks in configuration space toward a specified target structure.^28^ The remaining 200 pathways were produced in the absence of explicit solvent using the Framework Rigidity Optimized Dynamics Algorithm (FRODA),^36^ which samples conformations by enforcing stereochemical constraints and hydrophobic contacts in a geometric targeting framework.

**Figure 3.**
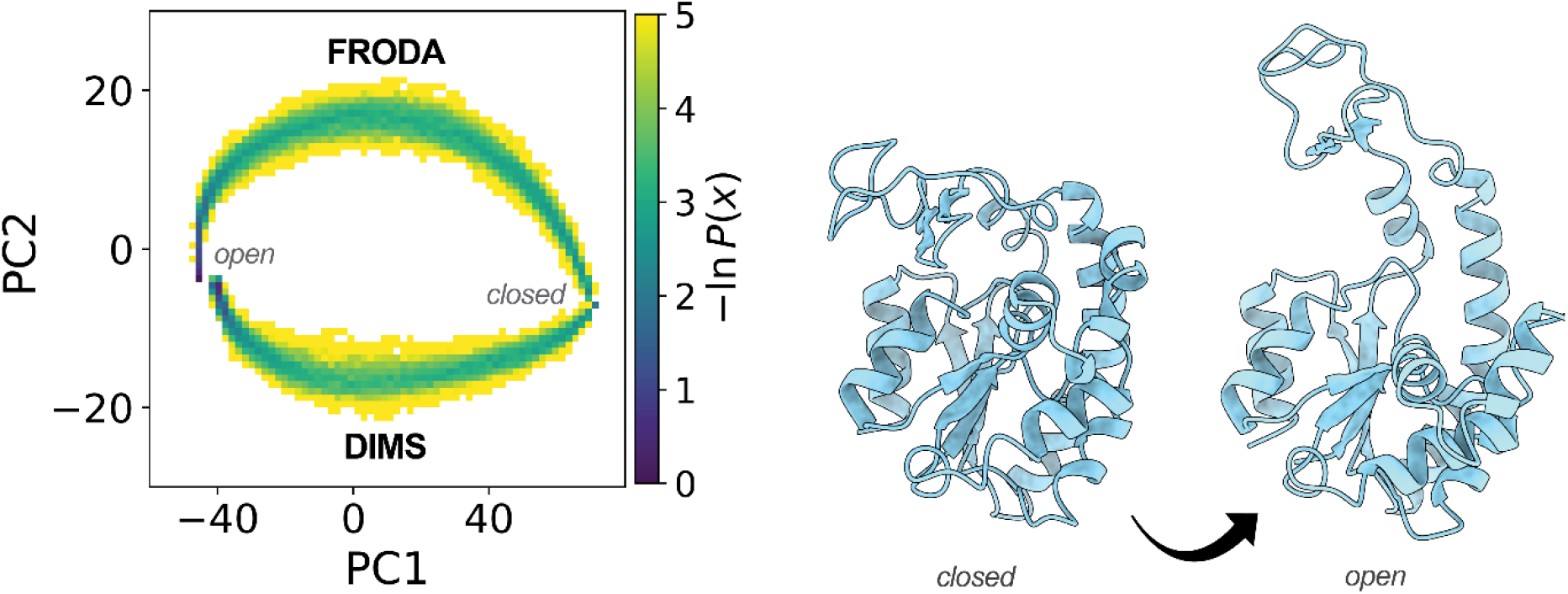
(Left) Probability density projected along the first two principal components (PC1 and PC2) capturing how the DIMS and FRODA sampling protocols explore distinct transition routes through the conformational space. (Right) Cartoon representations from the first DIMS trajectory showing the protein’s overall structure in its closed and open states.

The DIMS simulations employed the CHARMM22/CMAP protein force field^37^ with the ACS/ACE2 implicit solvent model and Langevin dynamics at 300 K. The FRODA pathways correspond to geometric trajectories between the same closed and open conformational states, providing a complementary ensemble of transition pathways with different underlying sampling assumptions.

### HIF-2_α_ System (HIF-2_α_)

The HIF-2α system is a protein-ligand complex consisting of the HIF-2α PAS-B domain as the protein receptor and a small-molecular ligand, THS-017 (Fig. 4). For this system, we used an ensemble of >4000 continuous ligand-unbinding pathways generated by Silvestrini et al.^30^ using WE simulations. The initial bound state of these simulations was generated by extracting heavy-atom coordinates from the corresponding crystal structure (PDB code: 3H7W).^38^ The protein was modeled using the Amber ff19SB force field and the ligand was modeled using compatible GAFF2 parameters. The protein-ligand complex was solvated in a truncated octahedral box of OPC water molecules^39^ and 17 mM NaCl ions using Li-Merz parameters that are compatible with the OPC water model.^40^ Dynamics were propagated with the Amber22 software package at a constant temperature of 25°C and constant pressure of 1 bar. The temperature was maintained using a weak Langevin thermostat with a collision frequency of 1 ps^-1^ and the pressure was maintained using a Monte Carlo barostat with pressure exchanges attempted every 0.2 ps.^41^

**Figure 4.**
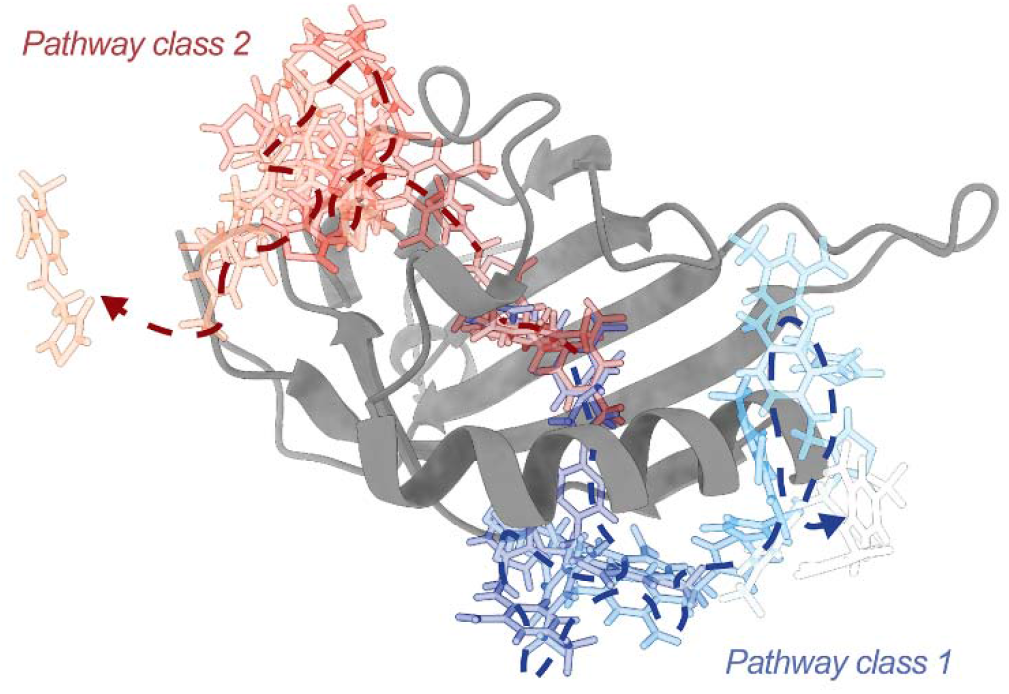
Structural representation of the protein-ligand complex featuring the HIF-2α PAS-B receptor (gray) and the small-molecule ligand THS-017, with overlaid ligand configurations tracing representative unbinding pathways as the molecule escapes from a completely buried internal cavity out into the bulk solvent.

The ensemble of unbinding pathways was generated from three independent WE simulations using the WESTPA 2.0 software.^32^ These WE simulations employed a three-dimensional progress coordinate consisting of (i) the solvent-accessible surface area (SASA) of the HIF-2⍰ internal cavity, (ii) a “ligand unbinding” RMSD, defined as the heavy-atom RMSD of the ligand after aligning on the receptor in the crystal structure of the ligand-receptor complex,^38^ and (iii) the ligand-receptor separation distance. This coordinate space was partitioned using a multi-stage, minimal adaptive binning scheme.^42^ To maintain a steady state, trajectories that reached a ligand-receptor separation distance of >10 Å were “recycled” by initiating a new trajectory with the same statistical weight from a randomly selected conformation within a pre-equilibrated ensemble of bound-state conformations. The initial bound-state was defined as having a ligand-receptor separation distance < 2.5 Å and ligand unbinding RMSD < 3 Å. To further accelerate simulation convergence, trajectories were reweighted using the weighted ensemble steady-state (WESS) method.^43^ All WE simulations employed a resampling time interval *τ* of 100 ps along with a target number of 8 trajectories per bin. The WE simulations yielded 35.1-41.5 µs of aggregate simulation time within 14-16 wall-clock days using 16 NVIDIA A100 GPUs in parallel.

## 4. RESULTS AND DISCUSSION

For each benchmark process, we compared PRISM using the three representative-set construction strategies (Options 1-3) against the original SHINE intra scheme. For PRISM, we examined *k* ∈ {6, 10,20, 50,80}, and for Option 3 we additionally considered *k*_final_ = *k* and *k*_final_ = 2*k*. This design allows us to assess (i) robustness to the granularity of medoid construction, (ii) sensitivity to the hierarchical refinement (for Option 3), and (iii) agreement with previously validated SHINE partitions.

### Alanine dipeptide

Alanine dipeptide served as a test case for recovering a small, sparsely populated pathway class from a dominant transition channel. Building on previous LPATH analysis, the least populated class was identified as a subset of trajectories labeled [25-27,33-47,51,68]. Our previous study demonstrated that intra- and semi-sum dissimilarities reliably recovered the core [33-47] subset, whereas inter, max, and classical Hausdorff distances often failed to capture this cluster structure.

Using PRISM, Options 1 and 2 reliably recovered this partition across all tested *k* values, consistently grouping the [33-47] trajectories within the minority pathway class (Fig. 5). At *k* = 80, all three options successfully recovered the target set [25-27,33-47,51,68] to a single cluster. This stability indicates that medoid-based abstraction preserves subtle mechanistic differences even when the number of representatives per pathway varies over an order of magnitude.

**Figure 5.**
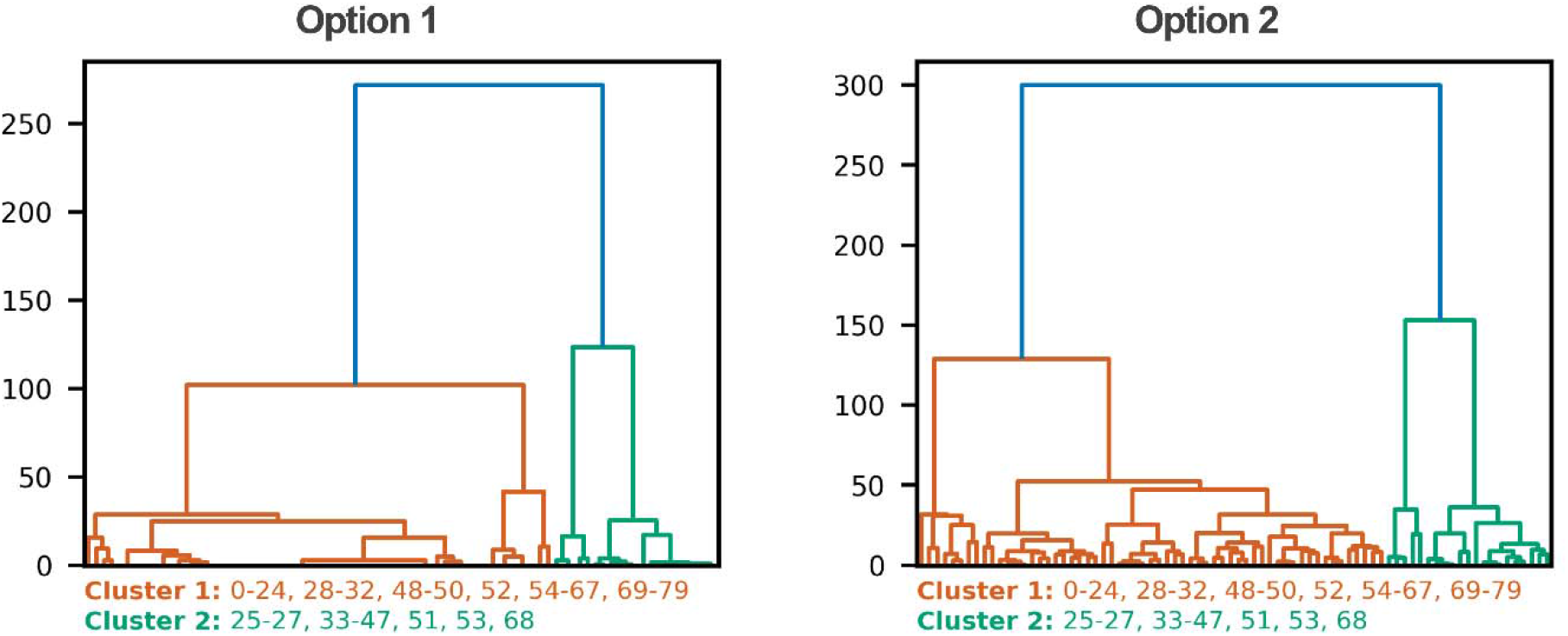
Alanine dipeptide pathway dendrograms for PRISM variants (Options 1, 2; *k* = 20). See Figs. S1-S6 for SHINE intra, Option 3, and additional *k* values.

In contrast, Option 3 exhibited greater sensitivity to *k* due to its two-stage refinement procedure. With *k*_final_ = *k*, the [33-47] trajectories were spuriously split into a third cluster only at *k* = 6, converging to the correct partition from *k* = 10 onward. With *k*_final_ = 2*k*, this sensitivity persisted longer where the previous observation appeared at *k* = 10, only correctly merging into the smaller cluster from *k* = 20 onward. This behavior reflects a resolution limitation of the two-stage procedure, when initial local clustering is too coarse, the pooled medoid set passed to the second stage lacks sufficient resolution to distinguish subtle geometric differences between pathway classes. As *k* increases, the medoids become more representative, allowing the two-stage construction to converge toward the robust behavior observed in Options 1 and 2.

### Adenylate Kinase

The AdK ensemble contains 400 closed-to-open transition pathways generated using two distinct sampling protocols: DIMS and FRODA. In the original SHINE analysis, the intra dissimilarity successfully separates these two methodological families. PRISM reproduces this separation across all three options and across all *k* values examined. The method consistently isolates the two pathway classes (Fig. 6). We also see that the separation occurs at well-defined linkage heights, indicating that the weighted average Hausdorff distance captures systematic geometric differences between the sampling mechanisms.

**Figure 6.**
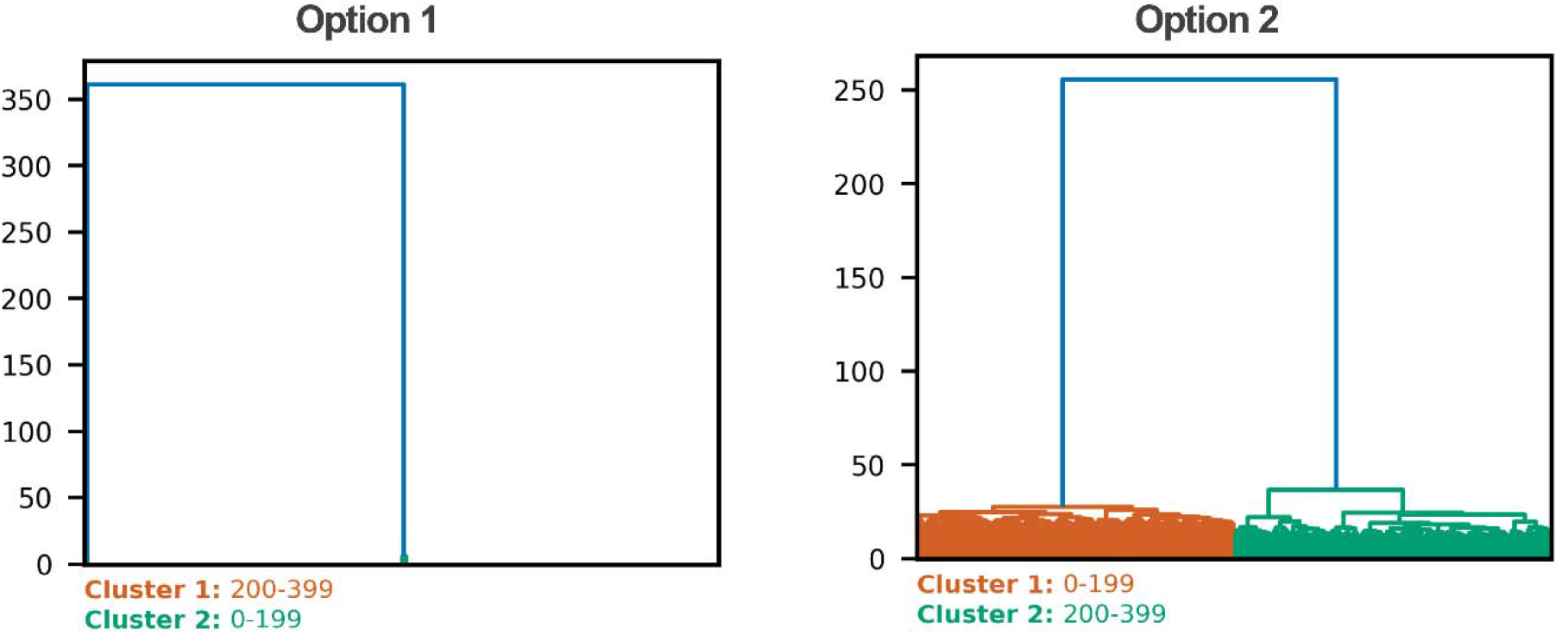
Adenylate kinase pathway dendrograms for PRISM variants (Options 1, 2; *k* = 20). See Figs. S7-S12 for SHINE intra, Option 3, and additional *k* values.

A qualitative difference between SHINE intra and PRISM emerges in the finer structure of the dendrogram near its base. In the intra scheme, merges occur progressively over a range of small, nonzero dissimilarities, creating a dense branching pattern. In contrast, PRISM produces tighter micro-clusters at very low linkage heights. This behavior arises because trajectories sharing identical or highly similar medoids have zero or near-zero average Hausdorff distances, leading to rapid consolidation of highly overlapping pathways. However, above this initial plateau the inter-pathway class merges occur at clearly separated heights, preserving the mechanistic interpretation. Thus, we show here that PRISM maintains agreement with SHINE at the level of dominant pathway families while offering a more compact structural representation.

### HIF-2⍰

The HIF-2⍰ ensemble is the largest and most heterogeneous dataset analyzed here with >4000 pathways. Using SHINE’s intra without sampling yields a roughly 52.0/48.0% population split between two dominant routes, whereas quota and diversity (50%) sampling shift the balance modestly toward 52.6/47.4% and 51.7/48.3%, respectively.

Across all three options, PRISM converges to a stable 53.4/46.4% partition by *k* = 20 (Fig. 7) and remains stable at all larger *k* values tested. Option 2 achieves this partition even at the smallest *k* = 6, reflecting that independent per-pathway clustering provides sufficient local coverage even with a few medoids. Option 1 requires a moderate number of global representatives to adequately cover the ensemble before the partition stabilizes, as a small global dictionary may fail to assign distinct representatives to structurally similar yet distinct pathways. Option 3 similarly converges by *k* = 20, consistent with the general observation that approaches relying on shared global representatives require a sufficient number of medoids to cleanly represent the diversity of pathways across the ensemble.

**Figure 7.**
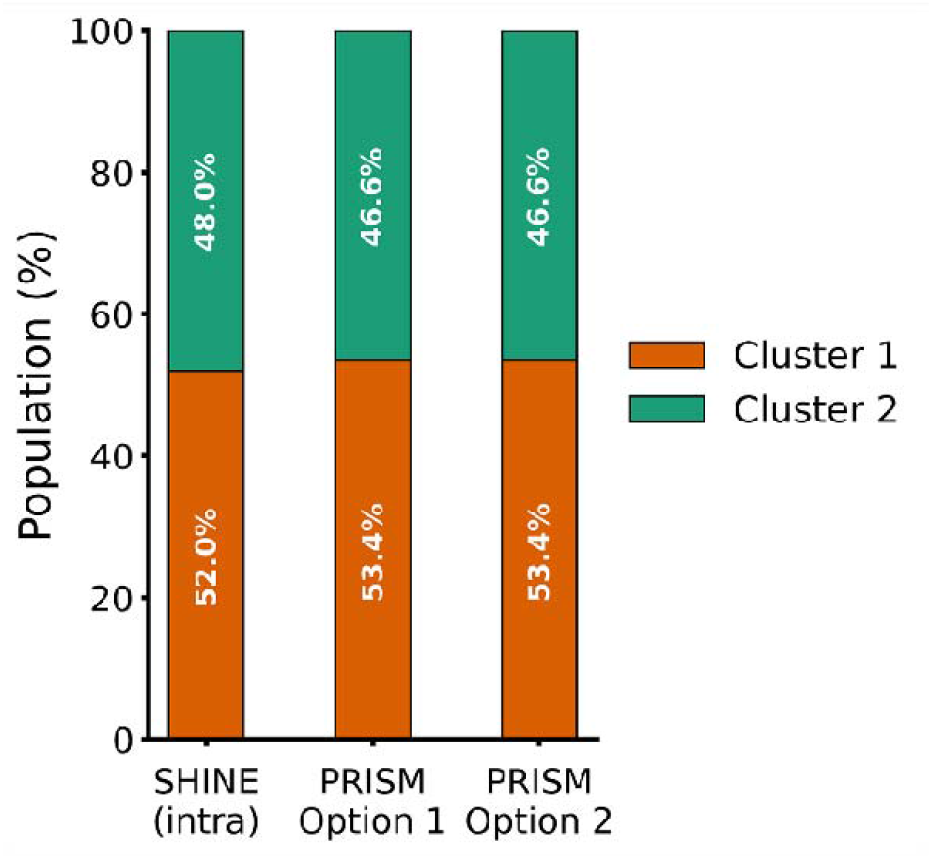
Population distribution of dominant ligand-unbinding pathways for the HIF-2⍰process, comparing the ensemble splits (Cluster 1 vs. Cluster 2) obtained using SHINE intra scheme (without sampling) and PRISM Options 1 and 2 with *k* = 20.

This behavior again highlights the importance of selecting sufficient representatives to accurately map each pathway, particularly for options that construct medoid sets from a shared global perspective, where a coarse dictionary risks collapsing structurally distinct pathways onto overlapping representatives. Unlike SHINE, PRISM achieves this population stability once a moderate *k* is reached, without requiring explicit frame sampling corrections. Detailed population percentages for SHINE and PRISM variants across the full range of tested *k* values are provided in the Supporting Information (Table S1).

### Computational Cost

Finally, we compared wall-clock runtimes for end-to-end clustering across the benchmark processes (Fig. 8); a comprehensive list of runtimes for SHINE and PRISM variants across full range of tested *k* is provided in the Supporting Information (Table S2). All timing benchmarks were performed on a single CPU core of an AMD Ryzen 5 5600G processor (3.9 GHz, 32 GB RAM) using a single Python process.

**Figure 8.**
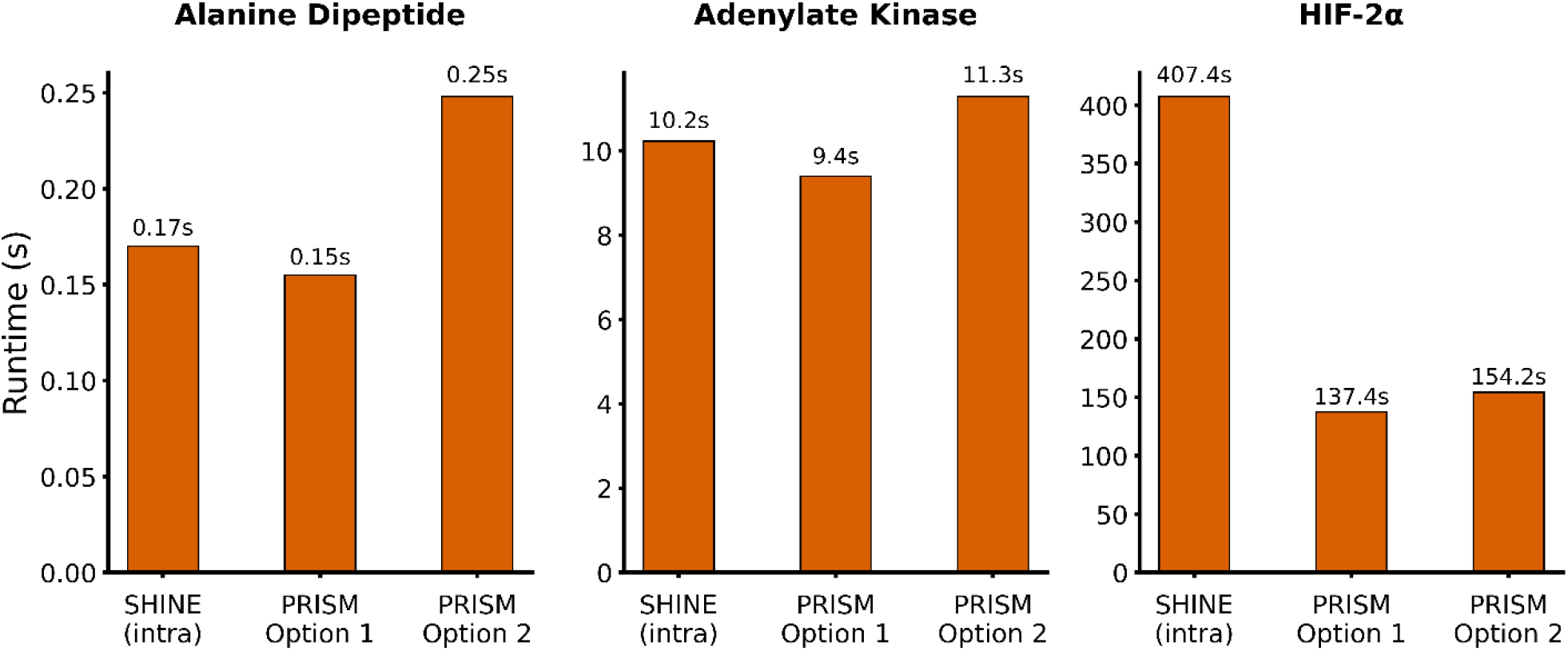
End-to-end clustering runtimes for the three benchmark processes using *k* = 20, comparing wall-clock times (in seconds) between SHINE intra scheme (without sampling) and PRISM Options 1 and 2 for (Left) alanine dipeptide, (Middle) adenylate kinase, and (Right) the HIF-2⍰system.

For the smallest system (ALA), all approaches completed in under one second, reflecting the limited size of the pathway ensemble and the negligible cost of distance construction at this scale; the only exception was intra with 50% diversity sampling, which introduced a measurable overhead even for the small system. For the AdK system, intra required approximately 10 s without sampling. While PRISM Options 1 and 3 achieved comparable runtimes, Option 2 exhibited a stronger dependence on representative-set size *k*, reflecting the tradeoff between a more detailed pathway summary and the cost of generating/processing it.

The most significant computational advantage of PRISM emerges for the largest ensemble (HIF-2⍰). In this case, intra without sampling required ~7 min, whereas diversity sampling, which explicitly searches for diverse, densely distributed frames, incurred substantially higher cost, increasing runtime to more than an hour. In contrast, PRISM maintained substantially shorter runtimes across all three options. Option 2 showed a modest increase at larger *k* (reaching up to ~5 min), consistent with its use of independent clustering per pathway that can increase computational overhead. Options 1 and 3 remain consistently faster, typically in the ~2-3 min range across the tested *k* values.

Overall, these results show that PRISM is competitive with, and often faster than, the original intra scheme, particularly when compared to computationally expensive diversity sampling strategies. This behavior aligns with the intended computational structure of PRISM: the modest overhead of computing medoids is offset by the efficiency of calculating pairwise-pathway dissimilarities between compact representative sets rather than between full frame ensembles.

### Practical Guidelines

The cluster number *k* controls the granularity with which trajectories are summarized before the pathway-pathway distances are computed. When *k* is small (e.g., 6-10) each trajectory can only be represented by a few medoids, which is often sufficient for systems dominated by a small number of well-separated conformational basins. This regime is also computationally efficient and typically recovers the dominant pathway families, but it can under-resolve subtle mechanistic distinctions when pathways differ primarily through localized excursions or intermediate states.

For most applications, we recommend an intermediate range of *k* (~20-50). In this regime, the number of representative medoids is large enough to capture finer structural variation within and across pathways while remaining compact relative to full frame ensembles. Empirically, this range tends to provide stable clustering results across processes, with minimal sensitivity to modest changes in *k*, and therefore serves as a practical default for routine analyses.

Larger *k* (e.g., *k* ≥ 80) can be advantageous for highly heterogeneous or flexible systems, where pathways may explore many distinct microstates and where coarse summaries may risk collapsing structurally meaningful variation. In practice, we observe that clustering results are often stable across *k* = 20-80, indicating that extreme precision in selecting *k* is generally unnecessary once representative sets are sufficiently expressive. A useful diagnostic is to monitor the population fractions of the dominant pathway clusters as functions of *k*. When these populations stabilize across successive *k* values, the representative sets are naturally adequate for robust clustering.

Moreover, choosing among PRISM’s three options primarily determines how local versus global structural organization is emphasized during medoid set construction. Option 1, which defines representatives in a shared space across the full ensemble, emphasizes global structural organization and can be advantageous when pathways repeatedly visit common basins, and one wishes to enforce a consistent global notion of “states” across trajectories. Option 2, which builds the representative sets locally for each pathway, is best suited for preserving pathway-specific structural states and is a natural default for mechanistic studies where subtle differences between individual trajectories are important.

Option 3 introduces a two-stage refinement procedure that balances local and global structure, which can be particularly useful for large and heterogeneous ensembles. In practice, Options 1 and 2 typically yield nearly identical dominant partitions in the benchmark processes considered here, while Option 3 can show mild sensitivity at very small *k* due to coarse initial local summaries. This sensitivity diminishes as *k* increases, and Option 3 converges to the same dominant clustering structure we see in the first two options.

## 5. CONCLUSIONS

We introduced PRISM, a state-aware framework for clustering molecular pathway ensembles that builds upon the conceptual framework of SHINE while adopting a distinct medoid-based representation strategy. By defining each pathway as a finite set of structural medoids obtained through *k*-means NANI clustering, PRISM decouples pathway comparison from full frame-level representations and instead operates on representative structural states. Pathway-pathway dissimilarities are then quantified using a weighted average Hausdorff distance, which measures the mean nearest-neighbor mismatch between representative sets in a geometrically interpretable manner.

To demonstrate the effectiveness of PRISM, we applied this clustering method to three benchmark processes: the C7_eq_ to C7_ax_ transition of alanine dipeptide, closed-to-open transition of adenylate kinase, and ligand-unbinding from the HIF-2α PAS domain. For all three processes, PRISM consistently recovers dominant mechanistic routes while exhibiting minimal sensitivity to the number of representatives *k* once moderate values are reached. In contrast to SHINE, PRISM does not require explicit frame sampling corrections to stabilize cluster populations, and it provides a more structurally interpretable representation of pathway families through the medoids. The method also scales naturally to large pathway ensembles and maintains deterministic behavior given a fixed construction strategy.

Moreover, several practical observations emerged from this study. First, clustering results are robust across a broad range of *k* (20-80), suggesting that PRISM does not require extensive parameter tuning. Second, the choice among the medoid-construction options primarily affects the balance between pathway-specific and globally shared structural features. Third, the weighted average Hausdorff formulation offers a stable geometric alternative to classical Hausdorff distance for pathway similarity by reducing sensitivity to structural outliers while preserving mechanistic resolution.

Overall, PRISM complements existing pathway-clustering approaches by linking trajectory-level similarity to compact, structurally interpretable representatives within a deterministic hierarchical framework. By extending SHINE’s philosophy into a representation-based regime, PRISM offers a practical balance of efficiency, robustness, and interpretability. This enables pathway ensembles to be organized into mechanistically meaningful families, providing stability against sampling noise and parameter sensitivity that is necessary for advancing analysis of biomolecular transitions.

## SUPPORTING INFORMATION

Alanine dipeptide pathway dendrograms for SHINE and PRISM variants (Figures S1–S6); adenylate kinase pathway dendrograms for SHINE and PRISM variants (Figures S7–S12); population distribution analysis for the HIF-2α system (Table S1); and wall-clock runtime comparisons for all benchmark processes (Table S2) (PDF).

## AUTHOR CONTRIBUTIONS

**JBWS**: Data curation; Formal analysis; Investigation; Software; Validation; Visualization; Writing. **JMGL**: Data curation; Formal analysis; Investigation; Software; Validation; Visualization; Writing. **LTC**: Formal analysis; Methodology; Conceptualization; Investigation; Software; Writing; Funding acquisition; Supervision; Resources. **RAMQ**: Formal analysis; Methodology; Conceptualization; Investigation; Software; Writing; Funding acquisition; Supervision; Resources.

## DATA AND SOFTWARE AVAILABILITY

The open-source PRISM code can be found here: https://github.com/mqcomplab/MDANCE.

## ACKNOWLEDGEMENTS

This research was supported by NIH grant R35GM150620 to RAMQ and NIH grant R01GM115805 to LTC.

## TOC Graphic

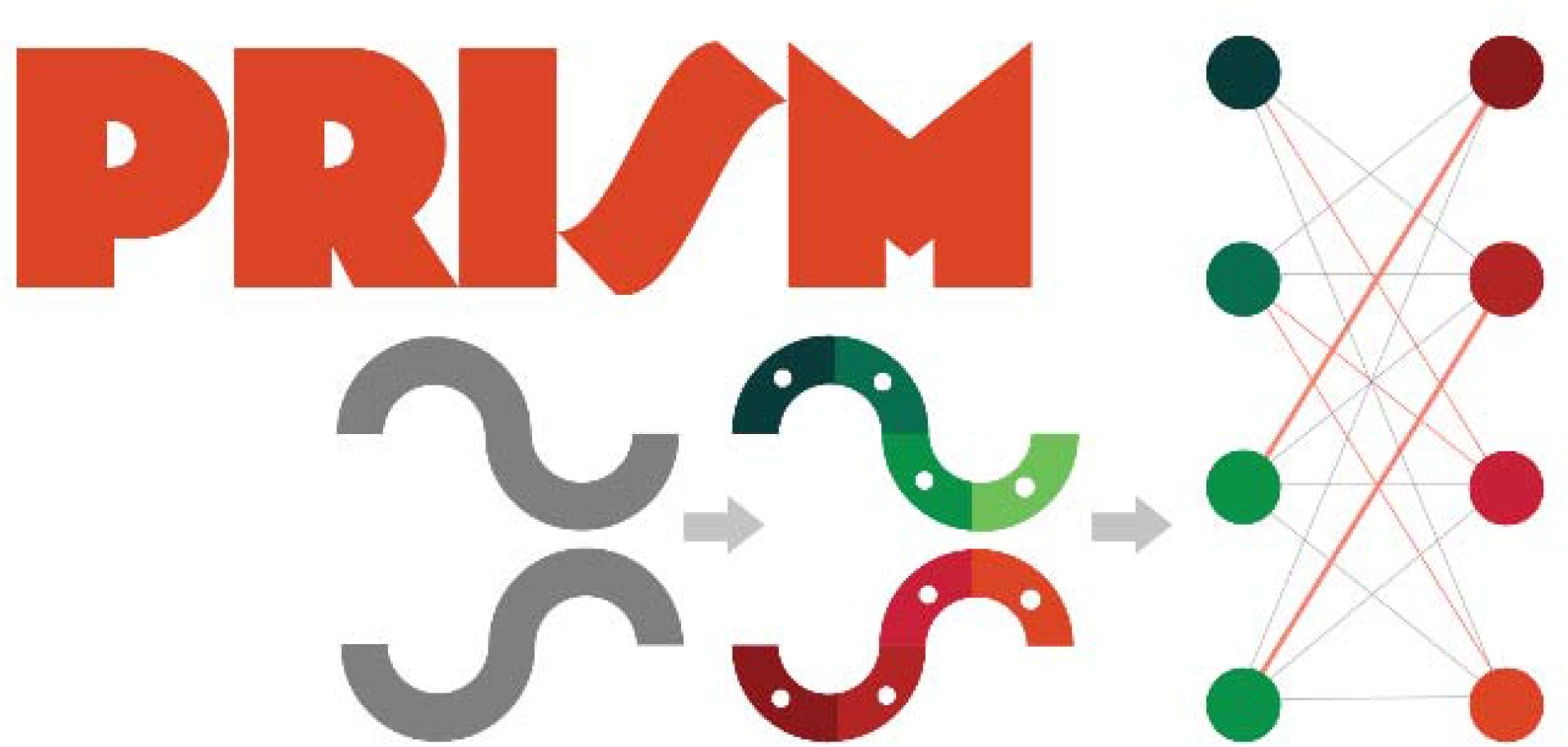

## References

(1) Kondratyuk, N.; Nikolskiy, V.; Pavlov, D.; Stegailov, V. GPU-Accelerated Molecular Dynamics: State-of-Art Software Performance and Porting from Nvidia CUDA to AMD HIP. The International Journal of High Performance Computing Applications 2021, 35 (4), 312–324. 10.1177/10943420211008288.

(2) Lee, T.-S.; Cerutti, D. S.; Mermelstein, D.; Lin, C.; LeGrand, S.; Giese, T. J.; Roitberg, A.; Case, D. A.; Walker, R. C.; York, D. M. GPU-Accelerated Molecular Dynamics and Free Energy Methods in Amber18: Performance Enhancements and New Features. J. Chem. Inf. Model. 2018, 58 (10), 2043–2050. 10.1021/acs.jcim.8b00462.

(3) Stone, J. E.; Hardy, D. J.; Ufimtsev, I. S.; Schulten, K. GPU-Accelerated Molecular Modeling Coming of Age. Journal of Molecular Graphics and Modelling 2010, 29 (2), 116–125. 10.1016/j.jmgm.2010.06.010.

(4) Voelz, V. A.; Pande, V. S.; Bowman, G. R. Folding@home: Achievements from over 20 Years of Citizen Science Herald the Exascale Era. Biophysical Journal 2023, 122 (14), 2852–2863. 10.1016/j.bpj.2023.03.028.

(5) Shaw, D. E.; Adams, P. J.; Azaria, A.; Bank, J. A.; Batson, B.; Bell, A.; Bergdorf, M.; Bhatt, J.; Butts, J. A.; Correia, T.; Dirks, R. M.; Dror, R. O.; Eastwood, M. P.; Edwards, B.; Even, A.; Feldmann, P.; Fenn, M.; Fenton, C. H.; Forte, A.; Gagliardo, J.; Gill, G.; Gorlatova, M.; Greskamp, B.; Grossman, J. P.; Gullingsrud, J.; Harper, A.; Hasenplaugh, W.; Heily, M.; Heshmat, B. C.; Hunt, J.; Ierardi, D. J.; Iserovich, L.; Jackson, B. L.; Johnson, N. P.; Kirk, M. M.; Klepeis, J. L.; Kuskin, J. S.; Mackenzie, K. M.; Mader, R. J.; McGowen, R.; McLaughlin, A.; Moraes, M. A.; Nasr, M. H.; Nociolo, L. J.; O’Donnell, L.; Parker, A.; Peticolas, J. L.; Pocina, G.; Predescu, C.; Quan, T.; Salmon, J. K.; Schwink, C.; Shim, K. S.; Siddique, N.; Spengler, J.; Szalay, T.; Tabladillo, R.; Tartler, R.; Taube, A. G.; Theobald, M.; Towles, B.; Vick, W.; Wang, S. C.; Wazlowski, M.; Weingarten, M. J.; Williams, J. M.; Yuh, K. A. Anton 3: Twenty Microseconds of Molecular Dynamics Simulation before Lunch. In Proceedings of the International Conference for High Performance Computing, Networking, Storage and Analysis; ACM: St. Louis Missouri, 2021; pp 1–11. 10.1145/3458817.3487397.

(6) Bolhuis, P. G.; Swenson, D. W. H. Transition Path Sampling as Markov Chain Monte Carlo of Trajectories: Recent Algorithms, Software, Applications, and Future Outlook. Advcd Theory and Sims 2021, 4 (4), 2000237. 10.1002/adts.202000237.

(7) Elber, R.; Fathizadeh, A.; Ma, P.; Wang, H. Modeling Molecular Kinetics with Milestoning. WIREs Comput Mol Sci 2021, 11 (4), e1512. 10.1002/wcms.1512.

(8) Chong, L. T.; Zuckerman, D. M. Weighted Ensemble Simulation: Advances in Methods, Software, and Applications. WIREs Comput Mol Sci 2025, 15 (6), e70055. 10.1002/wcms.70055.

(9) Chen, L.; Leung, J. M. G.; Zsigmond, K.; Chong, L. T.; Miranda-Quintana, R. A. SHINE: Deterministic Many-to-Many Clustering of Molecular Pathways. J. Chem. Inf. Model. 2025, 65 (10), 4775–4782. 10.1021/acs.jcim.5c00240.

(10) Seyler, S. L.; Kumar, A.; Thorpe, M. F.; Beckstein, O. Path Similarity Analysis: A Method for Quantifying Macromolecular Pathways. PLoS Comput Biol 2015, 11 (10), e1004568. 10.1371/journal.pcbi.1004568.

(11) Bogetti, A. T.; Leung, J. M. G.; Chong, L. T. LPATH: A Semiautomated Python Tool for Clustering Molecular Pathways. J. Chem. Inf. Model. 2023, 63 (24), 7610–7616. 10.1021/acs.jcim.3c01318.

(12) Georgouli, K.; Stephany, R. R.; Tempkin, J. O. B.; Santiago, C.; Aydin, F.; Heimann, M. A.; Pottier, L.; Zhang, X.; Carpenter, T. S.; Hsu, T.; Nissley, D. V.; Streitz, F. H.; Lightstone, F. C.; Ingolfsson, H. I.; Bremer, P.-T. Generating Protein Structures for Pathway Discovery Using Deep Learning. J. Chem. Theory Comput. 2024, 20 (20), 8795–8806. 10.1021/acs.jctc.4c00816.

(13) Gupta, A.; Ma, H.; Ramanathan, A.; Zerze, G. H. A Deep Learning-Driven Sampling Technique to Explore the Phase Space of an RNA Stem-Loop. J. Chem. Theory Comput. 2024, 20 (20), 9178–9189. 10.1021/acs.jctc.4c00669.

(14) Liu, B.; Boysen, J. G.; Unarta, I. C.; Du, X.; Li, Y.; Huang, X. Exploring Transition States of Protein Conformational Changes via Out-of-Distribution Detection in the Hyperspherical Latent Space. Nat Commun 2025, 16 (1), 349. 10.1038/s41467-024-55228-4.

(15) López-Pérez, K.; Kim, T. D.; Miranda-Quintana, R. A. iSIM: Instant Similarity. Digital Discovery 2024, 3 (6), 1160–1171. 10.1039/D4DD00041B.

(16) Miranda-Quintana, R. A.; Bajusz, D.; Rácz, A.; Héberger, K. Extended Similarity Indices: The Benefits of Comparing More than Two Objects Simultaneously. Part 1: Theory and Characteristics†. J Cheminform 2021, 13 (1), 32. 10.1186/s13321-021-00505-3.

(17) Miranda-Quintana, R. A.; Rácz, A.; Bajusz, D.; Héberger, K. Extended Similarity Indices: The Benefits of Comparing More than Two Objects Simultaneously. Part 2: Speed, Consistency, Diversity Selection. J Cheminform 2021, 13 (1), 33. 10.1186/s13321-021-00504-4.

(18) Chen, L.; Smith, M.; Roe, D. R.; Miranda-Quintana, R. A. Extended Quality (eQual): Radial Threshold Clustering Based on n-Ary Similarity. J. Chem. Inf. Model. 2025, 65 (10), 5062–5070. 10.1021/acs.jcim.4c02341.

(19) Chen, L.; Roe, D. R.; Miranda-Quintana, R. A. CADENCE: Clustering Algorithm─Density-Based Exploration and Novelty Clustering with Efficiency. J. Chem. Inf. Model. 2025, acs.jcim.5c00392. 10.1021/acs.jcim.5c00392.

(20) Chen, L.; Mondal, A.; Perez, A.; Miranda-Quintana, R. A. Protein Retrieval via Integrative Molecular Ensembles (PRIME) through Extended Similarity Indices. J. Chem. Theory Comput. 2024, 20 (14), 6303–6315. 10.1021/acs.jctc.4c00362.

(21) Miranda-Quintana, R. A.; Chen, L.; Santos, J. B. W. Molecular Dynamics Analysis with N-Ary Clustering Ensembles (MDANCE). https://github.com/mqcomplab/MDANCE.

(22) Chen, L.; Santos, J. B. W.; Gaza, J.; Perez, A.; Miranda-Quintana, R. A. Hierarchical Extended Linkage Method (HELM)’s Deep Dive into Hybrid Clustering Strategies. J. Chem. Inf. Model. 2025. 10.1021/acs.jcim.5c00539.

(23) Santos, J. B. W.; Chen, L.; Miranda-Quintana, R. A. Divide and Cluster: The DIVINE Framework for Deterministic Top-Down Analysis of Molecular Dynamics Trajectories. J. Chem. Inf. Model. 2026, acs.jcim.5c02740. 10.1021/acs.jcim.5c02740.

(24) Santos, J. B. W.; Chen, L.; Miranda-Quintana, R. A. mdBIRCH for Fast, Scalable, Online Clustering of Molecular Dynamics Trajectories. J. Chem. Theory Comput. 2026, 22 (8), 4026–4036. 10.1021/acs.jctc.6c00221.

(25) Noé, F.; Clementi, C. Collective Variables for the Study of Long-Time Kinetics from Molecular Trajectories: Theory and Methods. Current Opinion in Structural Biology 2017, 43, 141–147. 10.1016/j.sbi.2017.02.006.

(26) Chen, L.; Roe, D. R.; Kochert, M.; Simmerling, C.; Miranda-Quintana, R. A. K-Means NANI: An Improved Clustering Algorithm for Molecular Dynamics Simulations. J. Chem. Theory Comput. 2024, 20 (13), 5583–5597. 10.1021/acs.jctc.4c00308.

(27) Santos, J. B. W.; Chen, L.; Miranda-Quintana, R. A. Scaling *k*-Means for Multi-Million Frames: A Stratified NANI Approach for Large-Scale MD Simulations. J. Chem. Inf. Model. 2026, acs.jcim.5c02741. 10.1021/acs.jcim.5c02741.

(28) Seyler, S. L.; Beckstein, O. Sampling Large Conformational Transitions: Adenylate Kinase as a Testing Ground. Molecular Simulation 2014, 40 (10–11), 855–877. 10.1080/08927022.2014.919497.

(29) Beckstein, O.; Denning, E. J.; Perilla, J. R.; Woolf, T. B. Zipping and Unzipping of Adenylate Kinase: Atomistic Insights into the Ensemble of Open ↔ Closed Transitions. Journal of Molecular Biology 2009, 394 (1), 160–176. 10.1016/j.jmb.2009.09.009.

(30) Silvestrini, M. L.; Solazzo, R.; Boral, S.; Cocco, M. J.; Closson, J. D.; Masetti, M.; Gardner, K. H.; Chong, L. T. Gating Residues Govern Ligand Unbinding Kinetics from the Buried Cavity in HIF L2α PAS LB. Protein Science 2024, 33 (11), e5198. 10.1002/pro.5198.

(31) Fréchet, M. M. Sur quelques points du calcul fonctionnel. Rend. Circ. Matem. Palermo 1906, 22 (1), 1–72. 10.1007/BF03018603.

(32) Russo, J. D.; Zhang, S.; Leung, J. M. G.; Bogetti, A. T.; Thompson, J. P.; DeGrave, A. J.; Torrillo, P. A.; Pratt, A. J.; Wong, K. F.; Xia, J.; Copperman, J.; Adelman, J. L.; Zwier, M. C.; LeBard, D. N.; Zuckerman, D. M.; Chong, L. T. WESTPA 2.0: High-Performance Upgrades for Weighted Ensemble Simulations and Analysis of Longer-Timescale Applications. J. Chem. Theory Comput. 2022, 18 (2), 638–649. 10.1021/acs.jctc.1c01154.

(33) Maier, J. A.; Martinez, C.; Kasavajhala, K.; Wickstrom, L.; Hauser, K. E.; Simmerling, C. ff14SB: Improving the Accuracy of Protein Side Chain and Backbone Parameters from ff99SB. J. Chem. Theory Comput. 2015, 11 (8), 3696–3713. 10.1021/acs.jctc.5b00255.

(34) Onufriev, A. V.; Case, D. A. Generalized Born Implicit Solvent Models for Biomolecules. Annu. Rev. Biophys. 2019, 48 (1), 275–296. 10.1146/annurev-biophys-052118-115325.

(35) Brooks, B. R.; Brooks, C. L.; Mackerell, A. D.; Nilsson, L.; Petrella, R. J.; Roux, B.; Won, Y.; Archontis, G.; Bartels, C.; Boresch, S.; Caflisch, A.; Caves, L.; Cui, Q.; Dinner, A. R.; Feig, M.; Fischer, S.; Gao, J.; Hodoscek, M.; Im, W.; Kuczera, K.; Lazaridis, T.; Ma, J.; Ovchinnikov, V.; Paci, E.; Pastor, R. W.; Post, C. B.; Pu, J. Z.; Schaefer, M.; Tidor, B.; Venable, R. M.; Woodcock, H. L.; Wu, X.; Yang, W.; York, D. M.; Karplus, M. CHARMM: The Biomolecular Simulation Program. J Comput Chem 2009, 30 (10), 1545–1614. 10.1002/jcc.21287.

(36) Farrell, D. W.; Speranskiy, K.; Thorpe, M. F. Generating Stereochemically Acceptable Protein Pathways. Proteins 2010, 78 (14), 2908–2921. 10.1002/prot.22810.

(37) Buck, M.; Bouguet-Bonnet, S.; Pastor, R. W.; MacKerell, A. D. Importance of the CMAP Correction to the CHARMM22 Protein Force Field: Dynamics of Hen Lysozyme. Biophysical Journal 2006, 90 (4), L36–L38. 10.1529/biophysj.105.078154.

(38) Key, J.; Scheuermann, T. H.; Anderson, P. C.; Daggett, V.; Gardner, K. H. Principles of Ligand Binding within a Completely Buried Cavity in HIF2α PAS-B. J. Am. Chem. Soc. 2009, 131 (48), 17647–17654. 10.1021/ja9073062.

(39) Izadi, S.; Anandakrishnan, R.; Onufriev, A. V. Building Water Models: A Different Approach. J. Phys. Chem. Lett. 2014, 5 (21), 3863–3871. 10.1021/jz501780a.

(40) Sengupta, A.; Li, Z.; Song, L. F.; Li, P.; Merz, K. M. Parameterization of Monovalent Ions for the OPC3, OPC, TIP3P-FB, and TIP4P-FB Water Models. J. Chem. Inf. Model. 2021, 61 (2), 869–880. 10.1021/acs.jcim.0c01390.

(41) Allen, M. P.; Tildesley, D. J. Computer Simulation of Liquids: Michael P. Allen (Department of Physics, University of Warwick, UK, H.H. Wills Physics Laboratory, University of Bristol, UK), Dominic J. Tildesley (Centre Européen de Calcul Atomique et Moléculaire (CECAM), EPFL, Switzerland), Second edition.; Oxford University Press: Oxford, 2017.

(42) Torrillo, P. A.; Bogetti, A. T.; Chong, L. T. A Minimal, Adaptive Binning Scheme for Weighted Ensemble Simulations. J. Phys. Chem. A 2021, 125 (7), 1642–1649. 10.1021/acs.jpca.0c10724.

(43) Bhatt, D.; Zhang, B. W.; Zuckerman, D. M. Steady-State Simulations Using Weighted Ensemble Path Sampling. The Journal of Chemical Physics 2010, 133 (1), 014110. 10.1063/1.3456985.

