## Supplementary Information for "Pathway Representation via Intrinsic Structural Medoids (PRISM): A Structural Mapping Approach to Clustering Molecular Pathways"

| 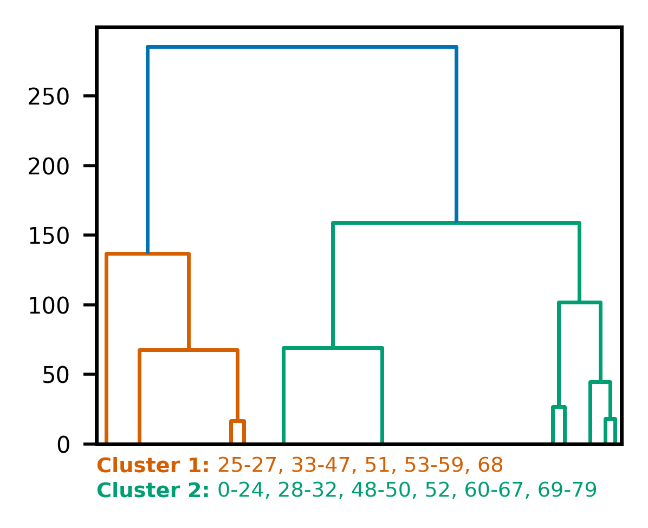  Option 1 | 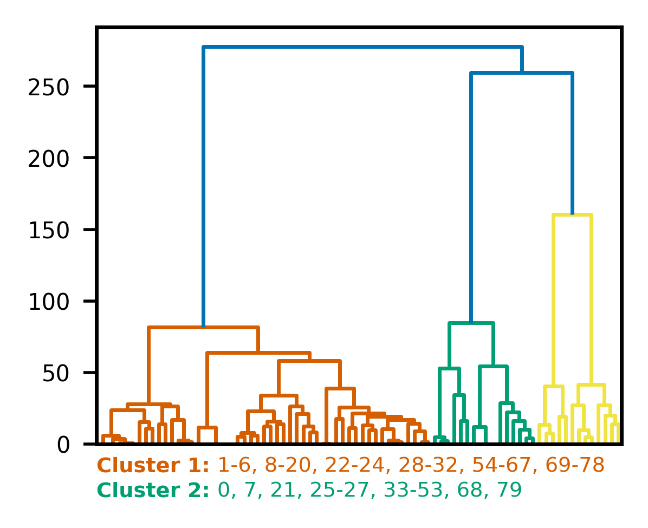  Option 2 |
| --- | --- |
| 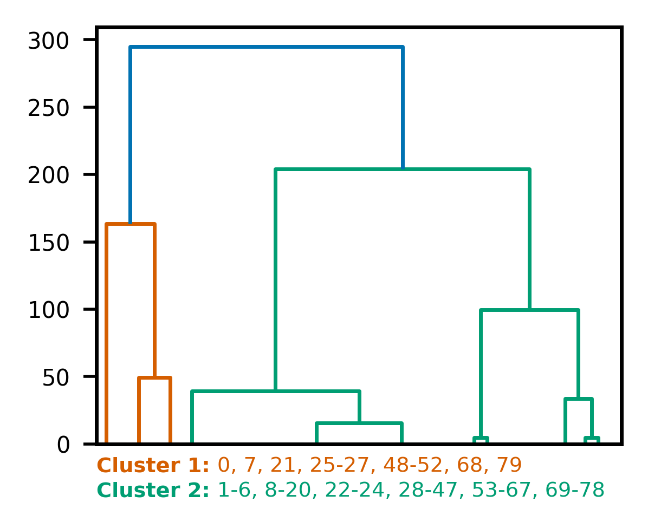  Option 3; *k*_final_ = *k* | 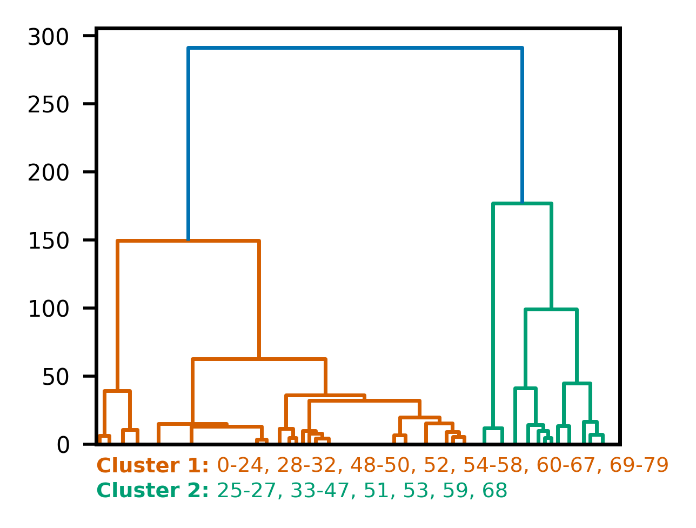  Option 3; *k*_final_ = 2*k* |

**Figure S1.** Alanine dipeptide pathway dendrograms for PRISM variants (Options 1, 2, 3; *k*=6).

| 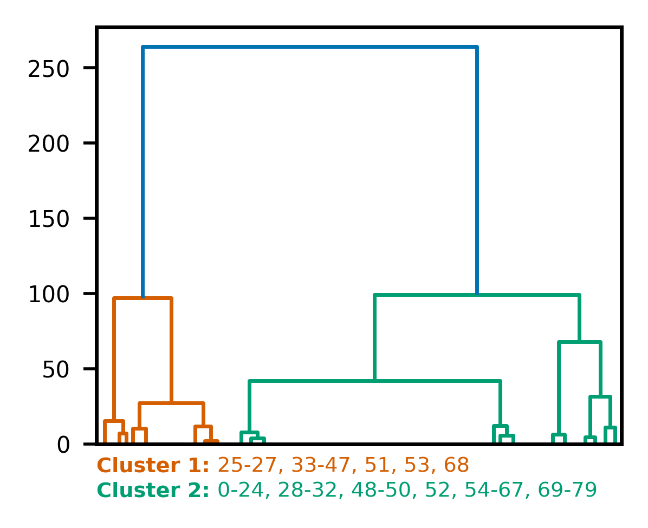  Option 1 | 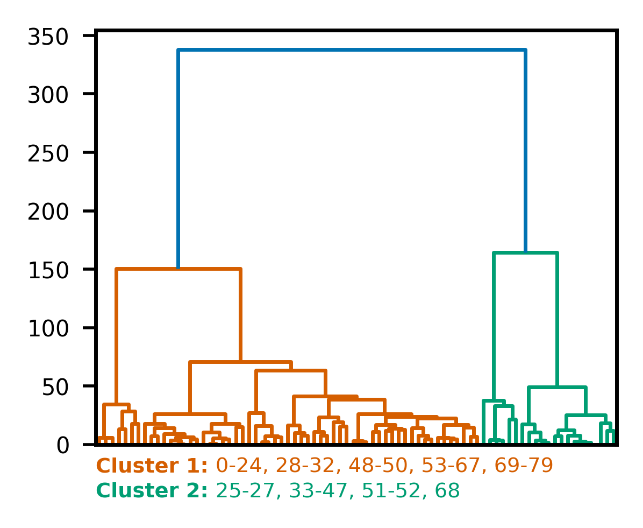  Option 2 |
| --- | --- |
| 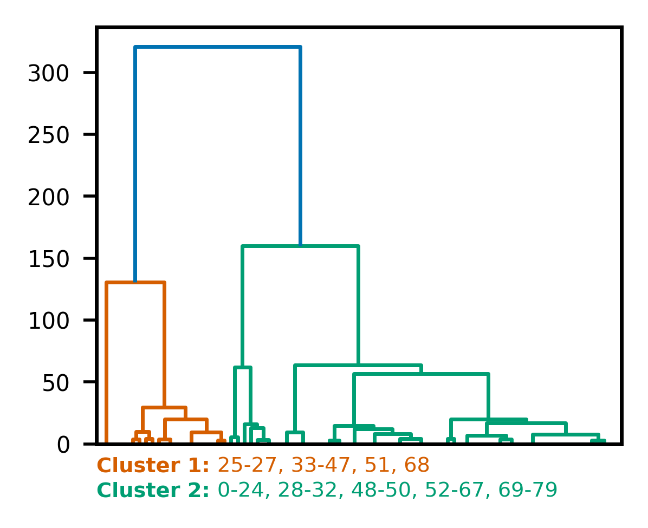  Option 3; *k*_final_ = *k* | 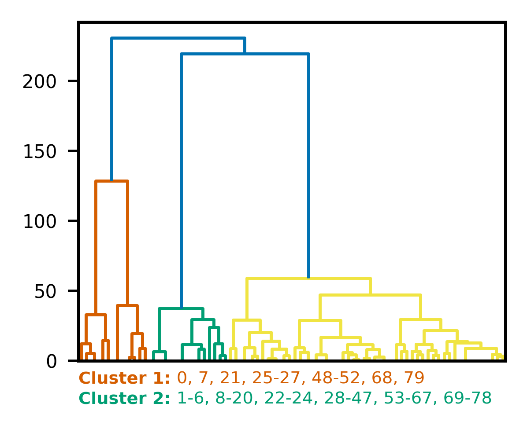  Option 3; *k*_final_ = 2*k* |

**Figure S2.** Alanine dipeptide pathway dendrograms for PRISM variants (Options 1, 2, 3; *k*=10).

| 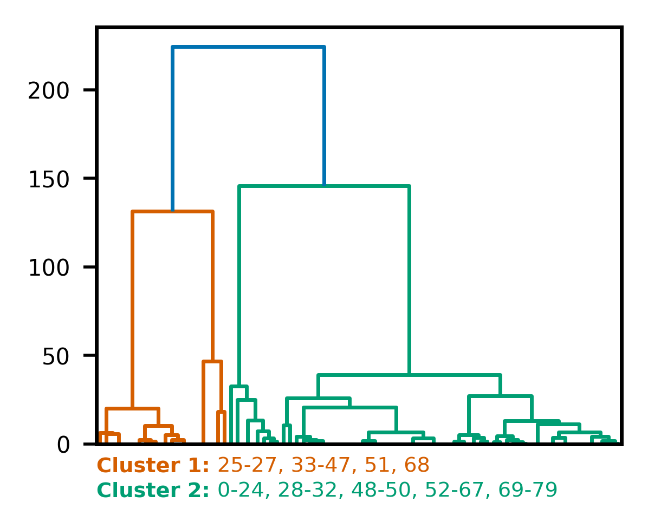  Option 3; *k*_final_ = *k* | 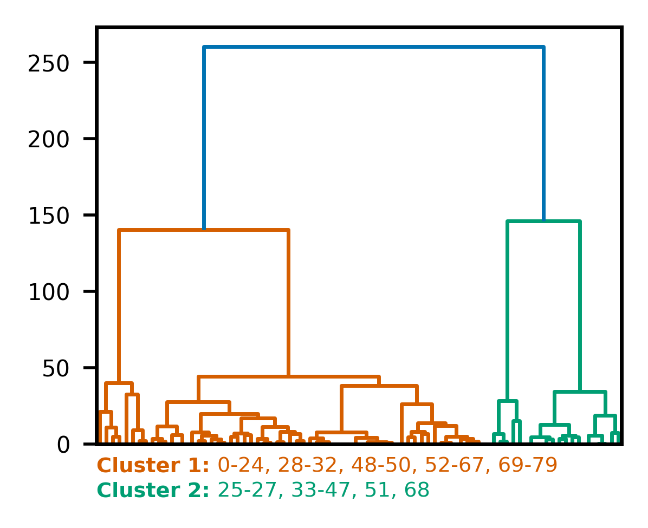  Option 3; *k*_final_ = 2*k* |
| --- | --- |

**Figure S3.** Alanine dipeptide pathway dendrograms for PRISM Option 3; *k*=20.

| 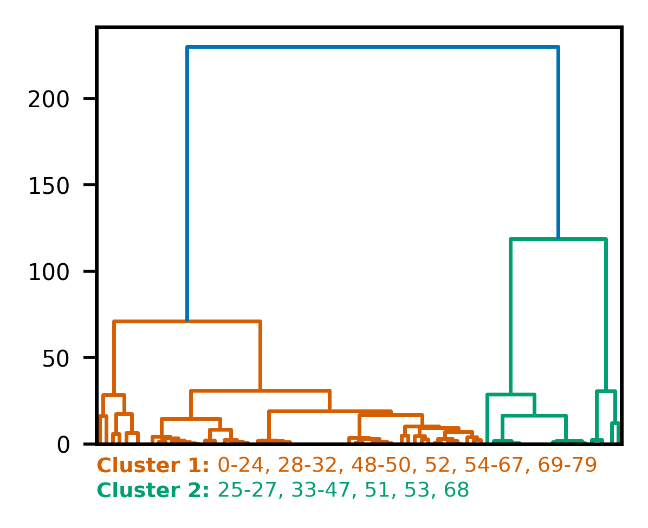  Option 1 | 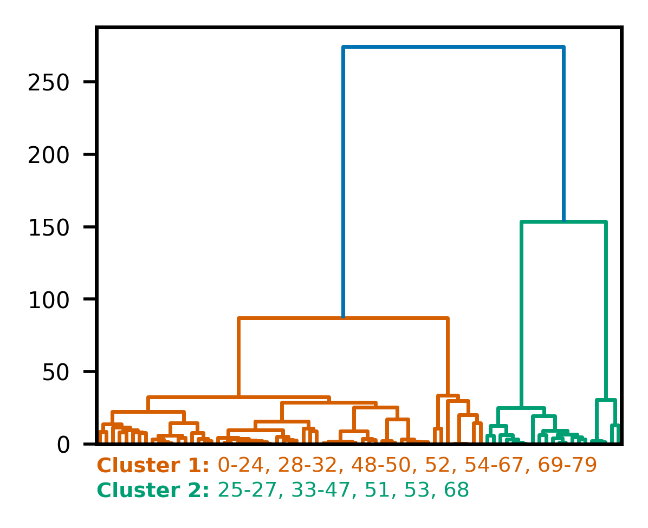  Option 2 |
| --- | --- |
| 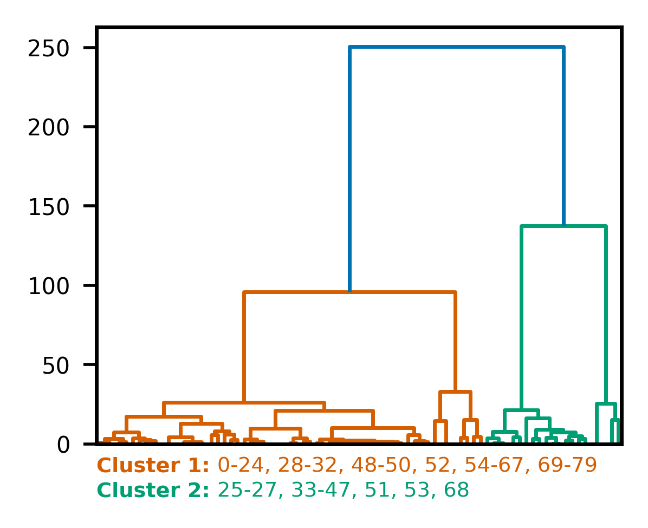  Option 3; *k*_final_ = *k* | 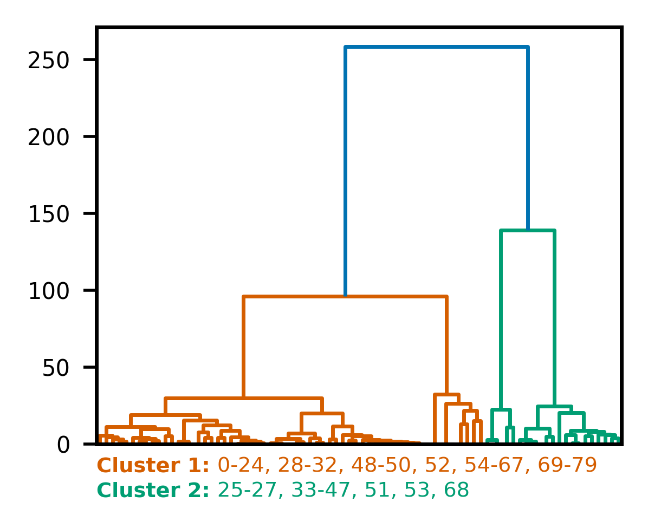  Option 3; *k*_final_ = 2*k* |

**Figure S4.** Alanine dipeptide pathway dendrograms for PRISM variants (Options 1, 2, 3; *k*=50).

| 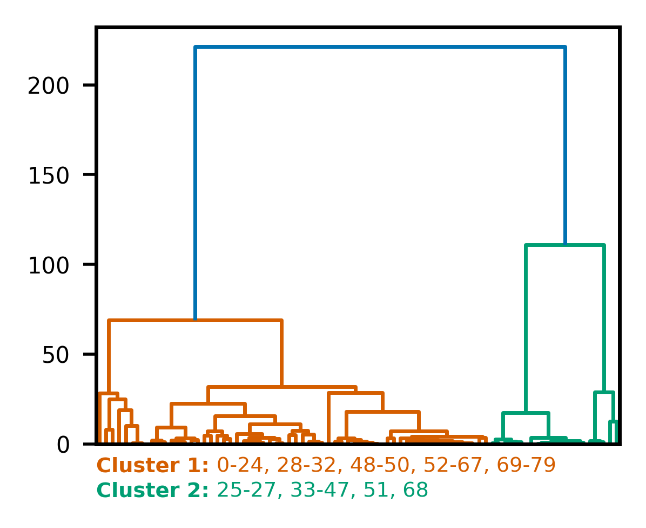  Option 1 | 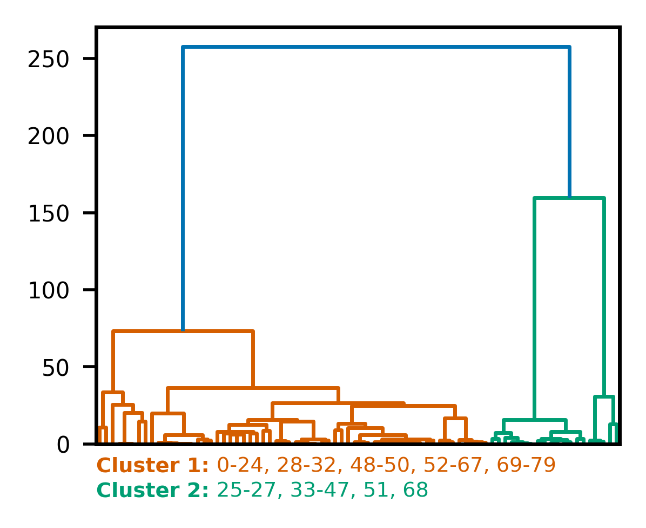  Option 2 |
| --- | --- |
| 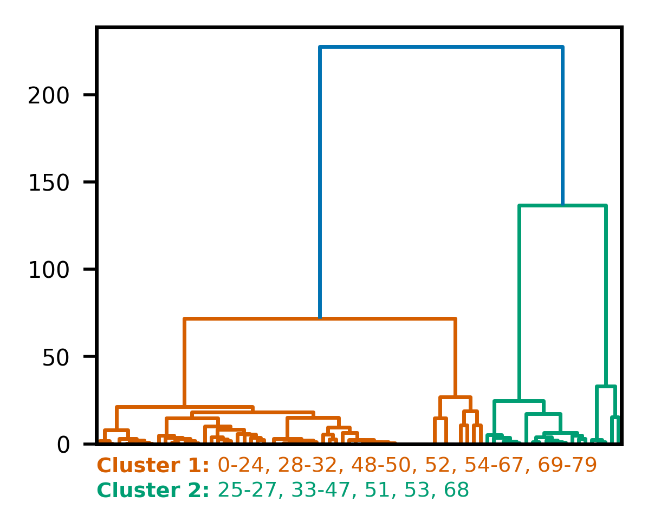  Option 3; $k_{\text{final}}= k$ | 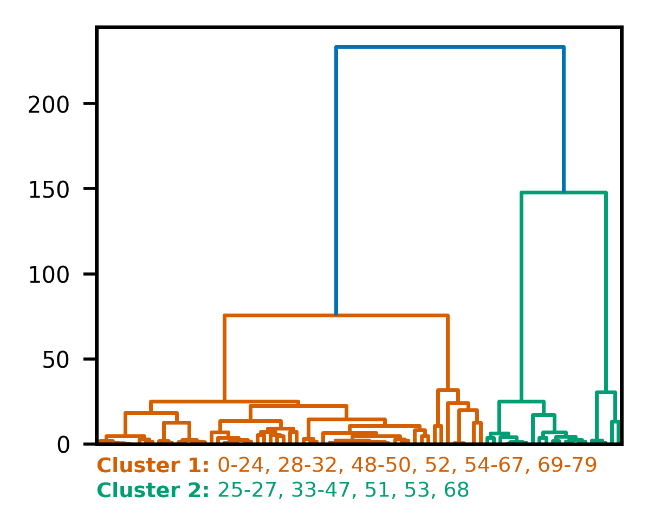  Option 3; $k_{\text{final}}= 2k$ |

**Figure S5.** Alanine dipeptide pathway dendrograms for PRISM variants (Options 1, 2, 3; *k*=80).

| 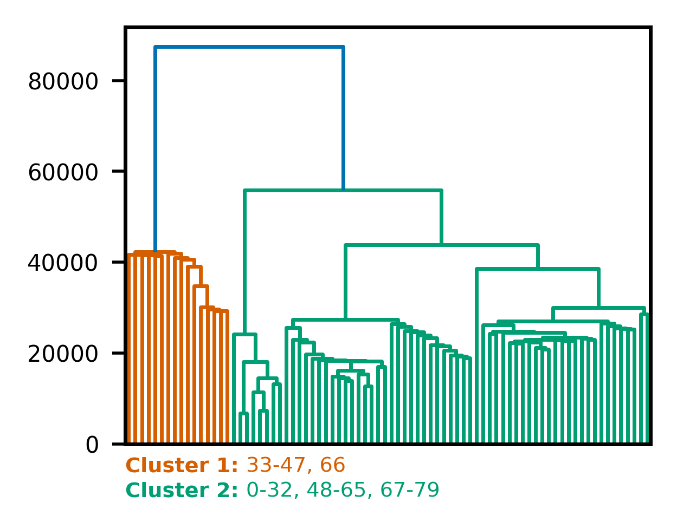  Intra; No Sampling | 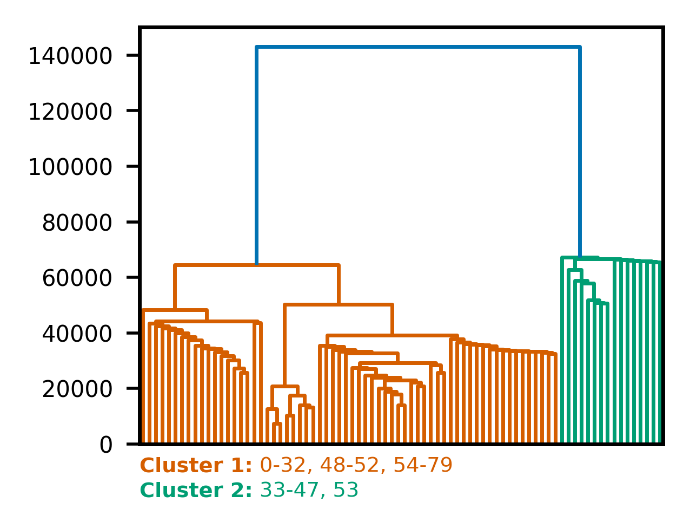  Intra; 50% Diversity Sampling |
| --- | --- |

**Figure S6.** Alanine dipeptide pathway dendrograms for SHINE intra (with and without sampling).

| 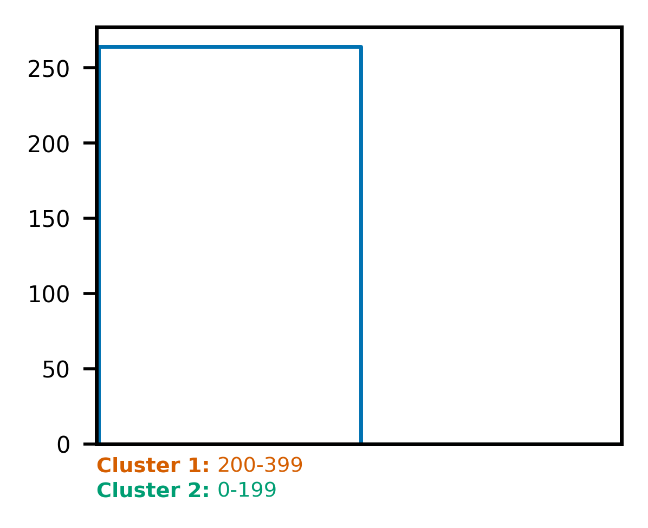  Option 1 | 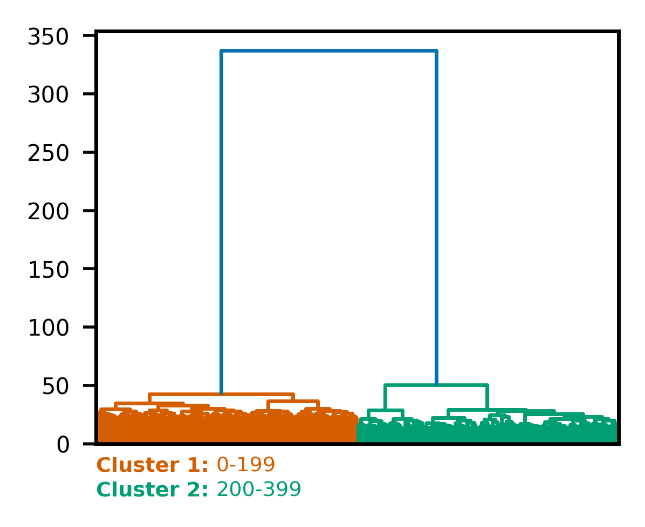  Option 2 |
| --- | --- |
| 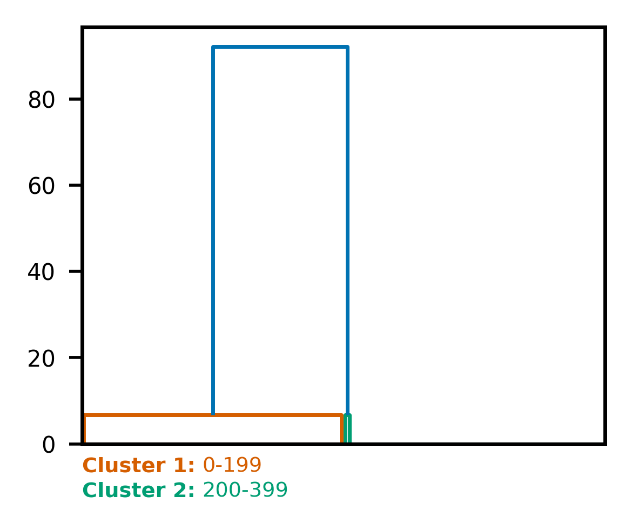  Option 3; *k*_final_ = *k* | 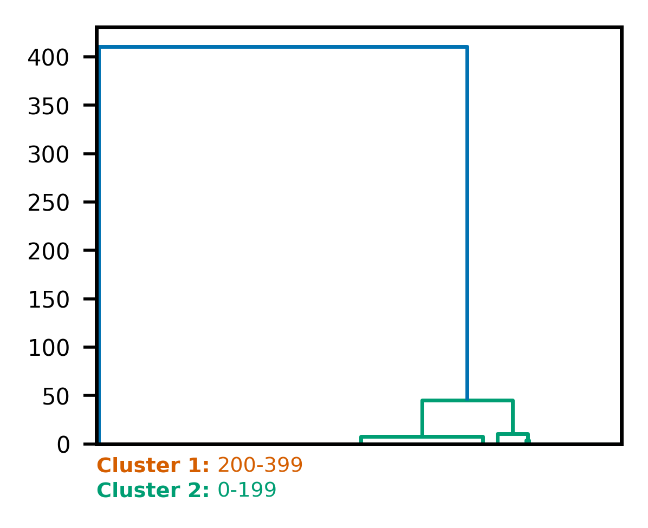  Option 3; *k*_final_ = 2*k* |

**Figure S7.** Adenylate kinase pathway dendrograms for PRISM variants (Options 1, 2, 3; *k*=6).

| 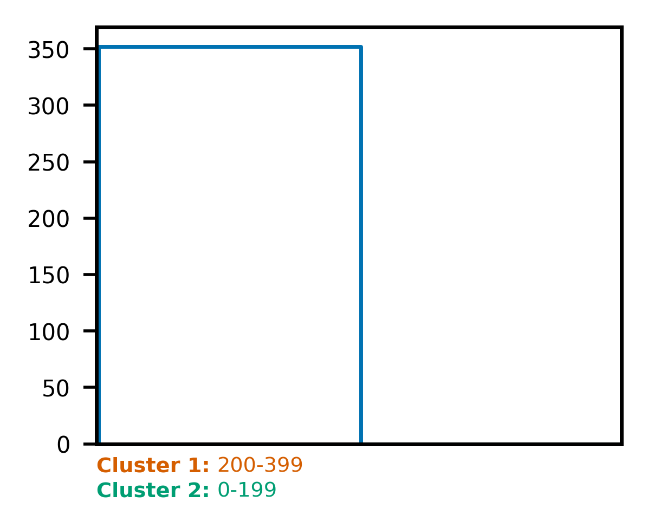  Option 1 | 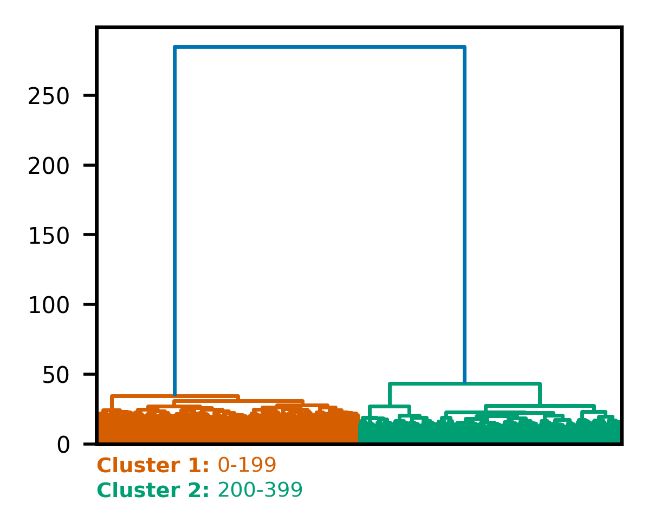  Option 2 |
| --- | --- |
| 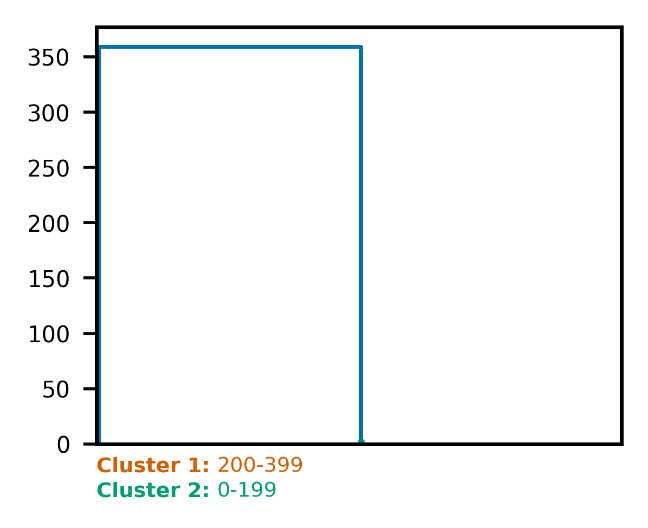  Option 3; *k*_final_ = *k* | 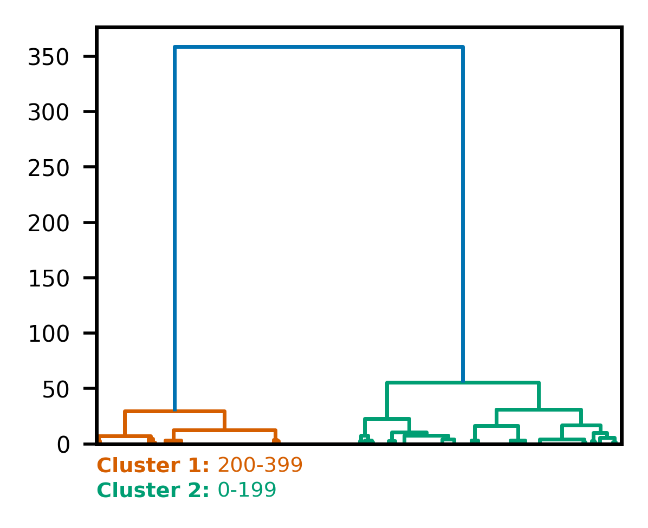  Option 3; *k*_final_ = 2*k* |

**Figure S8.** Adenylate kinase pathway dendrograms for PRISM variants (Options 1, 2, 3; *k*=10).

| 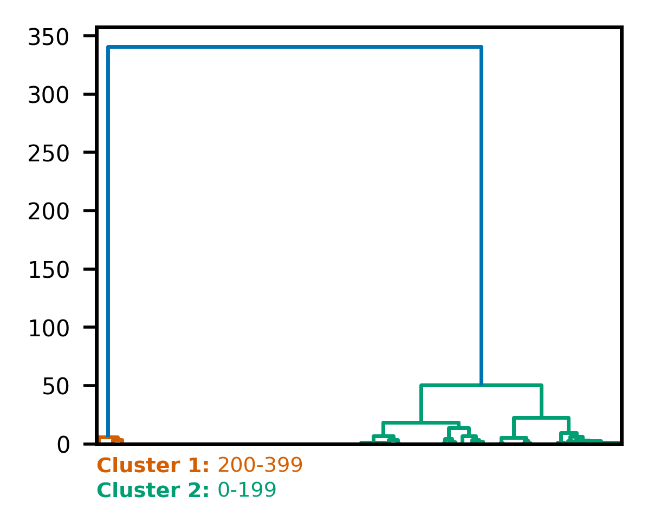  Option 3; *k*_final_ = *k* | 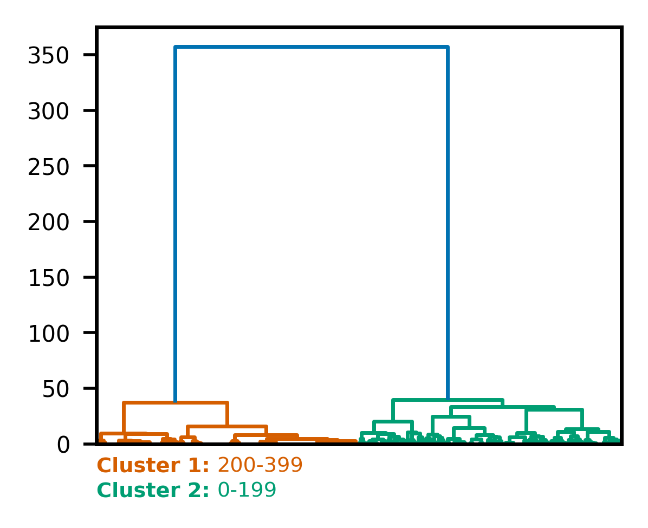  Option 3; *k*_final_ = 2*k* |
| --- | --- |

**Figure S9.** Adenylate kinase pathway dendrograms for PRISM Option 3; *k*=20.

| 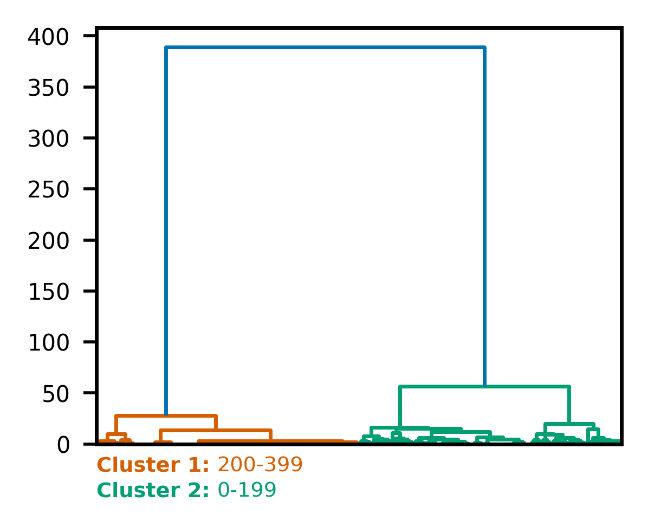  Option 1 | 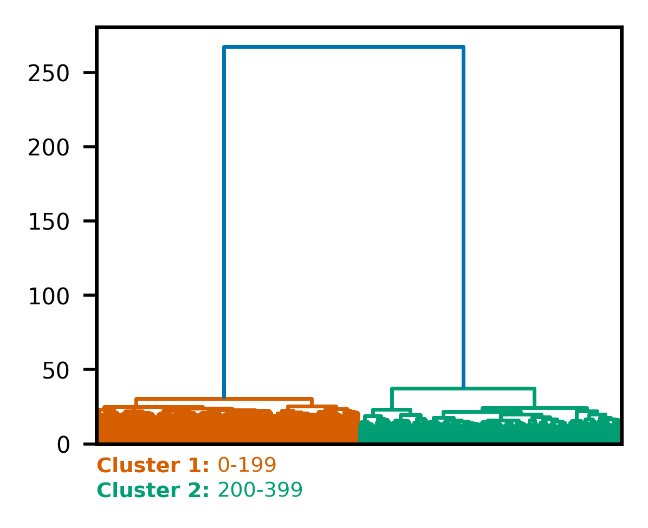  Option 2 |
| --- | --- |
| 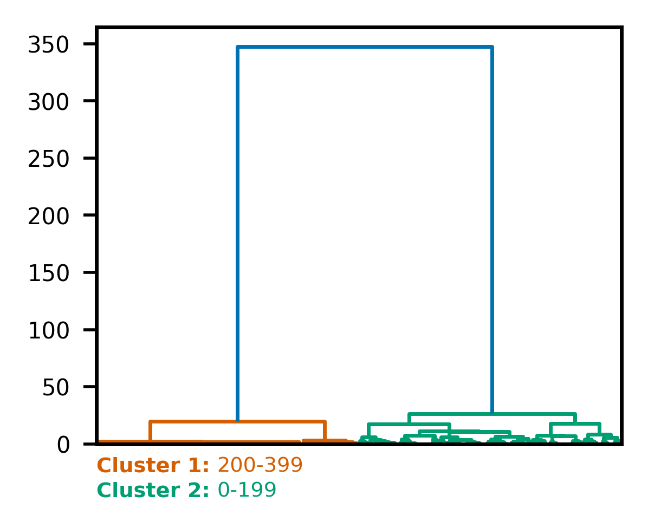  Option 3; *k*_final_ = *k* | 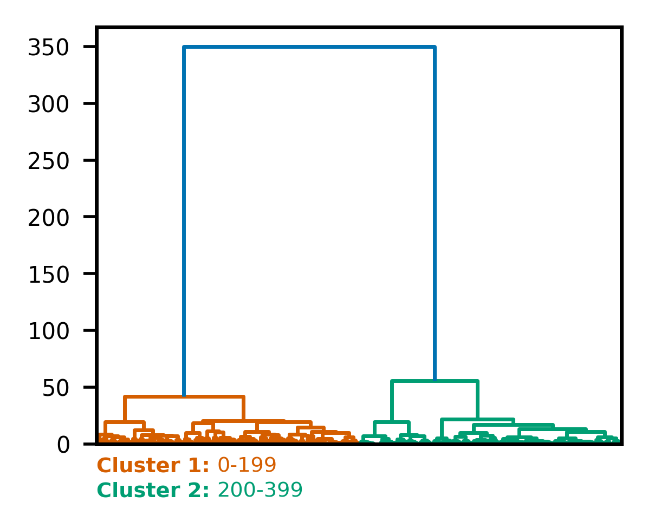  Option 3; *k*_final_ = 2*k* |

**Figure S10.** Adenylate kinase pathway dendrograms for PRISM variants (Options 1, 2, 3; *k*=50).

| 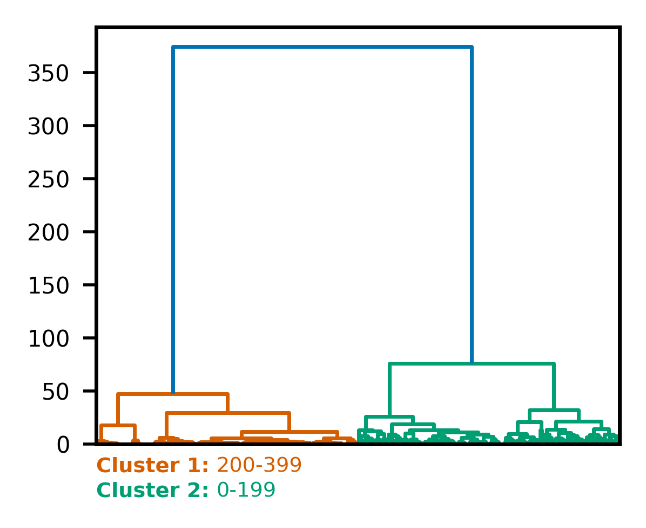  Option 1 | 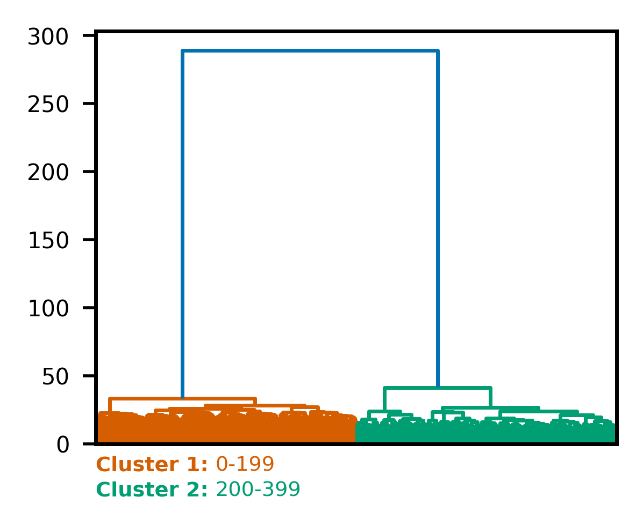  Option 2 |
| --- | --- |
| 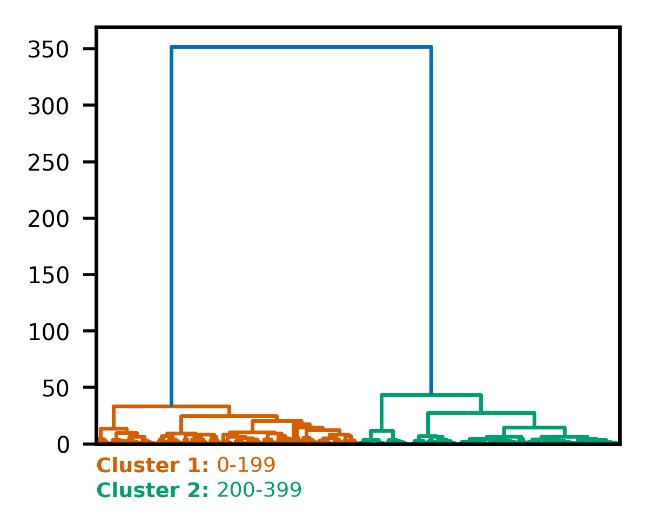  Option 3; $k_{\text{final}}= k$ | 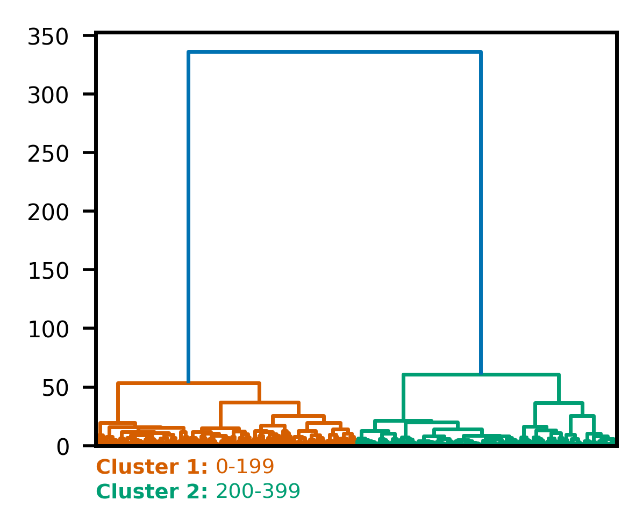  Option 3; $k_{\text{final}}= 2k$ |

**Figure S11.** Adenylate kinase pathway dendrograms for PRISM variants (Options 1, 2, 3; *k*=80).

| 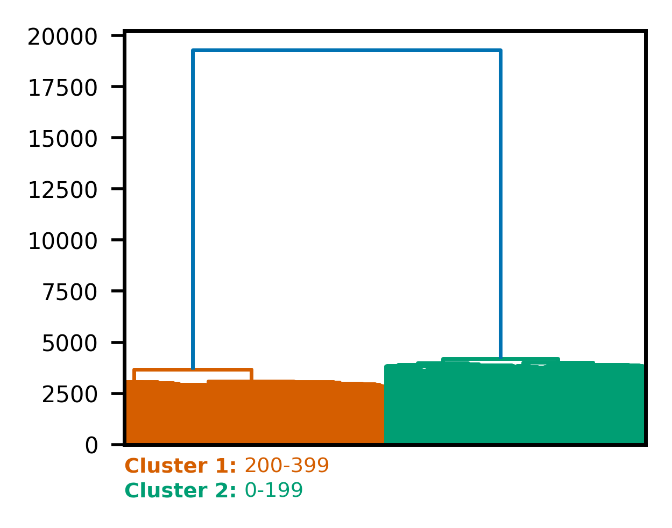  Intra; No Sampling | 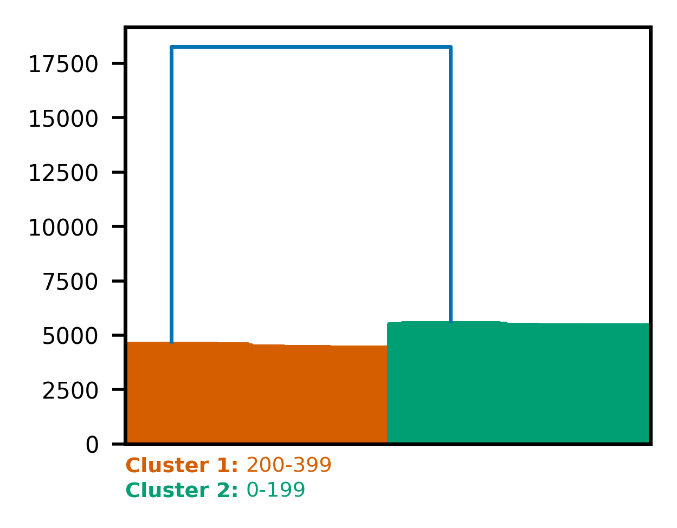  Intra; 50% Diversity Sampling |
| --- | --- |

**Figure S12.** Adenylate kinase pathway dendrograms for SHINE intra (with and without sampling).

**Table S1.** Population distribution analysis for the HIF-2⍺ system, comparing the population percentages of the two dominant clusters obtained using the SHINE intra scheme (with and without 50% sampling) and the three proposed PRISM variants at different cluster counts (*k*).

| **Method** | **Sampling** | ***k*** | **Cluster 1**  **Population (%)** | **Cluster 2**  **Population (%)** |
| --- | --- | --- | --- | --- |
| **SHINE**  **intra** | None | - | 52.0 | 48.0 |
|  | Quota | - | 52.6 | 47.4 |
|  | Diversity | - | 51.7 | 48.3 |
| **PRISM**  **Option 1** | None | 6 | 51.1 | 48.9 |
|  |  | 10 | 52.7 | 47.3 |
|  |  | 20 | 53.4 | 46.6 |
|  |  | 50 | 53.4 | 46.6 |
|  |  | 80 | 53.4 | 46.6 |
| **PRISM**  **Option 2** | None | 6 | 53.4 | 46.6 |
|  |  | 10 | 53.4 | 46.6 |
|  |  | 20 | 53.4 | 46.6 |
|  |  | 50 | 53.4 | 46.6 |
|  |  | 80 | 53.4 | 46.6 |
| **PRISM**  **Option 3**  $k_{\text{final}}=k$ | None | 6 | 53.8 | 46.2 |
|  |  | 10 | 53.4 | 46.6 |
|  |  | 20 | 53.4 | 46.6 |
|  |  | 50 | 53.4 | 46.6 |
|  |  | 80 | 53.4 | 46.6 |
| **PRISM**  **Option 3**  $k_{\text{final}}=2k$ | None | 6 | 53.4 | 46.6 |
|  |  | 10 | 53.5 | 46.5 |
|  |  | 20 | 53.4 | 46.6 |
|  |  | 50 | 53.4 | 46.6 |
|  |  | 80 | 53.4 | 46.6 |

**Table S2.** Comparison of clustering times (in seconds) across benchmark processes using the SHINE intra scheme (with and without 50% sampling) and the three proposed PRISM variants at different cluster counts (*k*). Runtime scaling benchmarks were performed on standard CPU hardware, and wall-clock times reflect end-to-end clustering runtimes.

| **Method** | **Sampling** | $\boldsymbol{k}$ | **ALA** | **AdK** | **HIF-2⍺** |
| --- | --- | --- | --- | --- | --- |
| **SHINE**  **intra** | None | - | 0.170 | 10.228 | 407.385 |
|  | Quota | - | 0.172 | 6.245 | 256.250 |
|  | Diversity | - | 4.862 | 24.678 | 3994.310 |
| **PRISM**  **Option 1** | None | 6 | 0.466 | 6.464 | 130.244 |
|  |  | 10 | 0.160 | 6.987 | 132.350 |
|  |  | 20 | 0.155 | 9.394 | 137.403 |
|  |  | 50 | 0.169 | 14.789 | 143.507 |
|  |  | 80 | 0.184 | 17.599 | 147.360 |
| **PRISM**  **Option 2** | None | 6 | 0.207 | 7.736 | 139.126 |
|  |  | 10 | 0.229 | 8.258 | 141.912 |
|  |  | 20 | 0.248 | 11.285 | 154.182 |
|  |  | 50 | 0.234 | 39.028 | 214.980 |
|  |  | 80 | 0.232 | 84.886 | 288.977 |
| **PRISM**  **Option 3**  $k_{\text{final}}=k$ | None | 6 | 0.217 | 7.029 | 136.678 |
|  |  | 10 | 0.227 | 7.130 | 135.875 |
|  |  | 20 | 0.227 | 7.774 | 137.203 |
|  |  | 50 | 0.250 | 12.004 | 149.234 |
|  |  | 80 | 0.258 | 14.600 | 153.326 |
| **PRISM**  **Option 3**  $k_{\text{final}}=2k$ | None | 6 | 0.216 | 7.433 | 135.153 |
|  |  | 10 | 0.231 | 7.732 | 141.489 |
|  |  | 20 | 0.240 | 8.773 | 141.890 |
|  |  | 50 | 0.274 | 12.906 | 148.802 |
|  |  | 80 | 0.278 | 15.254 | 155.526 |
